# Testing for genetic associations in arbitrarily structured populations

**DOI:** 10.1101/012682

**Authors:** Minsun Song, Wei Hao, John D. Storey

**Affiliations:** Lewis-Sigler Institute for Integrative Genomics, Princeton University, Princeton, NJ USA; Center for Statistics and Machine Learning, Princeton University, Princeton, NJ USA; Department of Molecular Biology, Princeton University, Princeton, NJ USA

## Abstract

We present a new statistical test of association between a trait and genetic markers, which we theoretically and practically prove to be robust to arbitrarily complex population structure. The statistical test involves a set of parameters that can be directly estimated from large-scale genotyping data, such as that measured in genome-wide association studies (GWAS). We also derive a new set of methodologies, called a genotype-conditional association test (GCAT), shown to provide accurate association tests in populations with complex structures, manifested in both the genetic and environmental contributions to the trait. We demonstrate the proposed method on a large simulation study and on the Northern Finland Birth Cohort study. In the Finland study, we identify several new significant loci that other methods do not detect. Our proposed framework provides a substantially different approach to the problem from existing methods, such as the linear mixed model and principal component approaches.

## INTRODUCTION

Performing genome-wide tests of association between a trait and genetic markers is one of the most important research efforts in modern genetics [1-3]. However, a major problem to overcome is how to test for associations in the presence of population structure [4]. Human populations are often structured in the sense that the genotype frequencies at a particular locus are not homogeneous throughout the population. Rather, there is heterogeneity in the genotype frequencies among individuals (correlated with variables such as geography or ancestry). At the same time, there may be other loci and non-genetic factors that also correlate with this genotype frequency heterogeneity, which in turn are correlated with the trait of interest. When this occurs, genetic markers become spuriously statistically associated with the trait of interest despite the fact that there is no biological connection.

The importance of addressing association testing in structured populations is evidenced by the existence of a large literature of methods proposed for this problem [5, 6]. The well-established methods all take a similar strategy in that the trait is modeled in terms of the genetic markers of interest, while attempting to adjust for genetic structure. Two popular approaches are to correct population structure by including principal components of genotypes as adjustment variables [7, 8] or by fitting a linear mixed effects model involving an estimated kinship or covariance matrix from the individuals’ genotypes [9, 10]. Previous work investigating the limitations of these two methods includes Wang, et al. (2013) [11]. These two approaches have been shown to be based on a common model that make differing assumptions about how the kinship or covariance matrices are utilized in the model [5]. This common model does not allow for nongenetic (e.g., environmental) contributions to the trait to be dependent with population structure. The linear mixed effects model requires that the genetic component is composed of small effects that additively are well-approximated by the Normal distribution. The model itself is therefore an approximation, and it is not yet possible to theoretically prove that a test based on this model is robust to structure for the more general class of relevant models that we investigate.

By taking a substantially different approach that essentially reverses the placement of the trait and genotype in the model, we formulate and provide a theoretical solution to the problem of association testing in structured populations for both quantitative and binary traits under general assumptions about the complexity of the population structure and its relationship to the trait through both genetic and non-genetic factors. This theoretical solution directly leads to a method for addressing the problem in practice that differs in key ways from the mixed model and principal component approaches. The method is straightforward: a model of structure is first estimated from the genotypes, and then a logistic regression is performed where the SNP genotypes are logistically regressed on the trait plus an adjustment based on the fitted structure model. The coefficient corresponding to the trait is then tested for statistical significance.

This association-testing framework is robust to general forms of population genetic structure, as well as to non-genetic effects that are dependent or correlated with population genetic structure (for example, lifestyle and environment may be correlated with ancestry) and with heteroskedasticity that is dependent on structure. We introduce an implementation of this test, called “genotype conditional association test” (GCAT). We show the proposed method corrects for structure on simulated data with a quantitative trait and compares favorably to existing methods. We also apply the method to the Northern Finland Birth Cohort data [12] and identify several new associated loci that have not been identified by existing methods. For example, the proposed method is the only one to identify a SNP (rs2814982) associated with height, which we note is linked to another SNP (rs2814993) that has been associated with skeletal frame size [13]. We discuss the advantages and disadvantages of the proposed framework with existing approaches, and we conclude that the proposed framework will be useful in future studies as sample sizes and the complexity of structure increase.

## RESULTS

### Motivation and Rationale of the Proposed Association Test

We have developed a new statistical framework to test for associations between genetic markers and a trait (either binary or quantitative) in the presence of population structure. Importantly, this test also allows for non-genetic factors correlated with population structure to contribute to the trait. Full mathematical and algorithmic details of the proposed framework are given in Online Methods and Supplementary Note, including mathematical proofs that the proposed framework is immune to general forms of population structure and non-genetic effects that are correlated with this structure.

One way in which spurious associations occur in the presence of population structure is that SNPs become correlated with each other when structure is not taken into account. Therefore, if a SNP is causal for the trait of interest, then any other SNP correlated with this causal SNP may also show an association. For SNPs in linkage disequilibrium due to their physical proximity with the causal SNP, one expects these to be associated with the trait regardless of structure. (This is a key property underlying the GWAS strategy.) However, in the presence of structure, there may be many unlinked SNPs that also yield spurious associations with the trait due to the fact that structure induces correlations of these SNPs with the causal SNP. Indeed, one of the early methods for detecting structure in association studies was to show that many randomly chosen, unlinked SNPs show associations to the trait [4]. This source of confounding is typically the main focus of association tests designed for structured populations.

Another key issue that is less often considered is the fact that lifestyle, environment, and other non-genetic factors that contribute to trait variation are often correlated with population structure (Fig. 1a). (This is the case because non-genetic factors are often strongly related to geography and ancestry, for example.) An association test that is immune to population structure should also be immune to the non-genetic effects that are confounded with structure. The framework we developed therefore also accounts for non-genetic factors that influence the trait and are correlated with population genetic structure.

**Figure 1.**
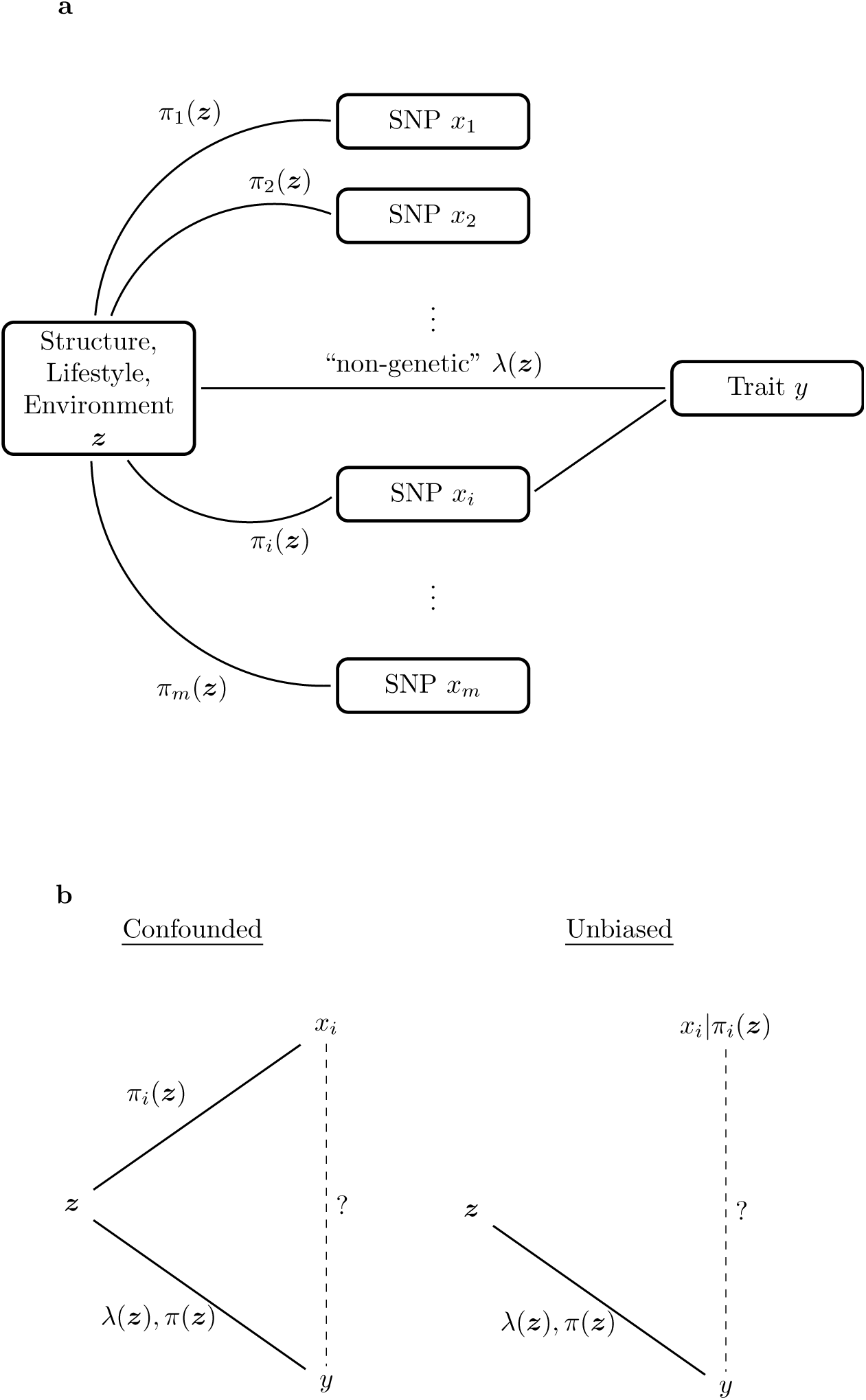
Rationale for the proposed test of association. (a) A graphical model describing population structure and its effects on a trait of interest. Population structure is captured by a common latent variable ***z*** among a set of loci *x_i_*(*i* = 1,2, …, *m*), via the allele frequencies *π_i_*(***z***). When one locus has a causal effect on the trait, this induces spurious associations with other loci affected by population structure. At the same time, population structure may be correlated with lifestyle and environment as these are all possibly related to ancestry and geography. (b) Accounting for confounding due to latent population structure. Left panel: A test for association between the *i*^th^ SNP *x_i_* and trait *y* without taking into account ***z*** will produce a spurious association due to the fact that both *x_i_* and *y* are confounded with ***z***. Right panel: A test for association between *x_i_*|*π_i_*(***z***) and *y* will be an unbiased because conditioning on *π_i_*(***z***) breaks the relationship between ***z*** and *x_i_*.

The rationale for the proposed test is schematized in Fig. 1. In the presence of population structure and non-genetic factors, the SNP *x_i_* and the trait y become spuriously associated due to a confounding latent variable ***z***. This latent variable is interpreted as capturing information on individual-specific allele frequencies (e.g., capturing the effect of population structure), lifestyle, and environment, all of which may be interdependent and play a determining role in the trait. The problem is that we cannot directly observe ***z*** and we would like to avoid making assumptions about its mathematical form. If we can successfully construct either *x_i_*|***z*** (the distribution of *x_i_* conditional on ***z***) or *y*|***z***, then it is possible to perform a test of association between *x_i_* and *y* that is immune to the effects of ***z***. Unbiased association tests can occur between *x_i_*|***z*** and *y*, between *x_i_* and *y*|***z***, or between *x_i_*|***z*** and *y*|***z***.

The linear mixed model and principal components approaches can be interpreted as attempts to estimate a model of *y*|***z***. This requires additional assumptions about non-genetic and genetic effects, and their relationship to ***z***, specifically that there is no relationship between structure and non-genetic effects in the trait model (Online Methods, Supplementary Note, and ref. [5]).

Due to the massive number of SNPs that have been measured in GWAS, trying to instead construct *x_i_*|***z*** is appealing since information can be leveraged among many simultaneously genotyped SNPs to infer accurate models of population structure. (This information is readily visualized in principal components constructed from the genotypes.) Our approach is therefore to carry out an association test between *x_i_*|***z*** and *y* by specifically testing whether there is equality or not between Pr(*x_i_*|*y*, ***z***) and Pr(*x_i_*|***z***) (Fig. 1b). If Pr(*x_i_*|*y*, ***z***) = Pr(*x_i_*|***z***) then there is no association between the SNP *x_i_* and the trait *y*; if Pr(*x_i_*|*y*, ***z***) ≠ Pr(*x_i_*|***z***), then there is an association. The test of association defined in this manner is immune to population structure and correlated non-genetic effects because we have taken into account ***z***.

One remaining problem is that we cannot observe ***z***. However, it is feasible to estimate a model of *x_i_*|***z*** since this conditional distribution is characterized with a proper model of population genetic structure. If we let *π_i_*(***z***) be the allele frequency of an individual with latent variable ***z***, then the distribution of *x_i_*|***z*** is equivalent to that of *x_i_*|π_i_(***z***) under our assumptions. The test of association then becomes a test of whether Pr(*x_i_*|*y*, *π_i_*(***z***)) = Pr(*x_i_*|*π_i_*(***z***)) or not. Because the latent variable ***z*** is not directly utilized in the test, we are able to account non-genetic effects correlated with population structure without the need to directly observe or model them.

We have recently developed a framework, called logistic factor analysis (LFA), that flexibly models and estimates *x_i_*|*π_i_*(***z***) [14]. (Note that other models of *x_i_*|*π_i_*(***z***) exist and can be utilized within the framework; see Online Methods and Supplementary Note.) LFA forms a linear latent variable model of logit(*π_i_*(***z***), where 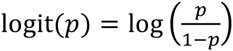 for 0 < *p* < 1. The number of latent variables, denoted by *d*, is chosen in Hao et al. [14] to best satisfy the model assumption that *x_i_*|*π_i_*(***z***) ~ Binomial(2, *π_i_*(***z***)).

To carry out the proposed association test, we perform a logistic regression of the SNP genotypes *x_i_* on the trait *y* plus the transformed individual-specific allele frequencies, logit(*π_i_*(***z***)). This reversal of the genotype and trait in the model fitting is an example of a technique from statistics called “inverse regression” and is distinct from the linear mixed model and principal components approaches. The trait models we assume are very general, so in practice the key assumption that needs to be verified is whether the model of *x_i_*|*π_i_*(***z***) is valid. This can be straightforwardly performed on the genotype data, and this process does not involve the trait variable or its underlying model, in particular its non-genetic effects that cannot be directly observed.

### Simulation Studies

We performed an extensive set of simulations to demonstrate that the proposed test is robust to population structure and to assess its power to detect true associations (full technical details in Online Methods and Supplementary Note). We compared the proposed test to its Oracle version (where model (3) from Online Methods and test-statistic (6) from Supplementary Note are used with the true individual-specific allele frequency values imputed). We also included in the simulation studies three important and popular methods: (i) the method of adjusting the trait and genotypes by principal components computed from the full set of genotypes [8] and (ii) two implementations of the linear mixed effects model (LMM) approach [9, 10], specifically EMMAX by Kang et al. (2010) [10] and GEMMA by Zhou and Stephens (2012) [15]. The methods are abbreviated as “PCA,” LMM-EMMAX,” and “LMM-GEMMA.”

For each of 33 simulation configurations, we simulated and analyzed 100 GWAS data sets from a quantitative trait model (equation (1) from Online Methods), for a grand total of 3300 simulated data sets. Each simulation scenario involved *m* = 100,000 simulated SNPs on *n* individuals, where *n* ranged from 940 to 5000 depending on the scenario. For a given simulated study, we therefore obtained a set of 100,000 p-values, one per SNP. So-called “spurious associations” occur when the p-values corresponding to null (non-associated) SNPs are artificially small. For a given p-value threshold *t*, we expect there to be *m*_0_×*t* false positives among the *m*_0_ p-values corresponding to null SNPs, where *m*_0_ = 100,000 − 10 in our case. At the same time, we can calculate the observed number of false positive simply by counting how many of the null SNP p-values are less than or equal to *t*. The excess observed false positives are spurious associations. A method properly accounts for structure when the average difference is zero. The best one can do on a study-by-study basis is captured by the Oracle method, which according to our theory is immune to structure and provides the correct null distribution.

Fig. 2 shows the excess in observed false positives vs. the expected number of false positives for the Oracle, GCAT (proposed), PCA, and both implementations of LMM under five configurations of structure for a quantitative trait variation apportionment corresponding to genetic=5%, non-genetic=5%, and noise=90%. It can be seen that the proposed GCAT method performs similarly to the Oracle test, whereas PCA tends to suffer from an excess of spurious associations.

**Figure 2.**
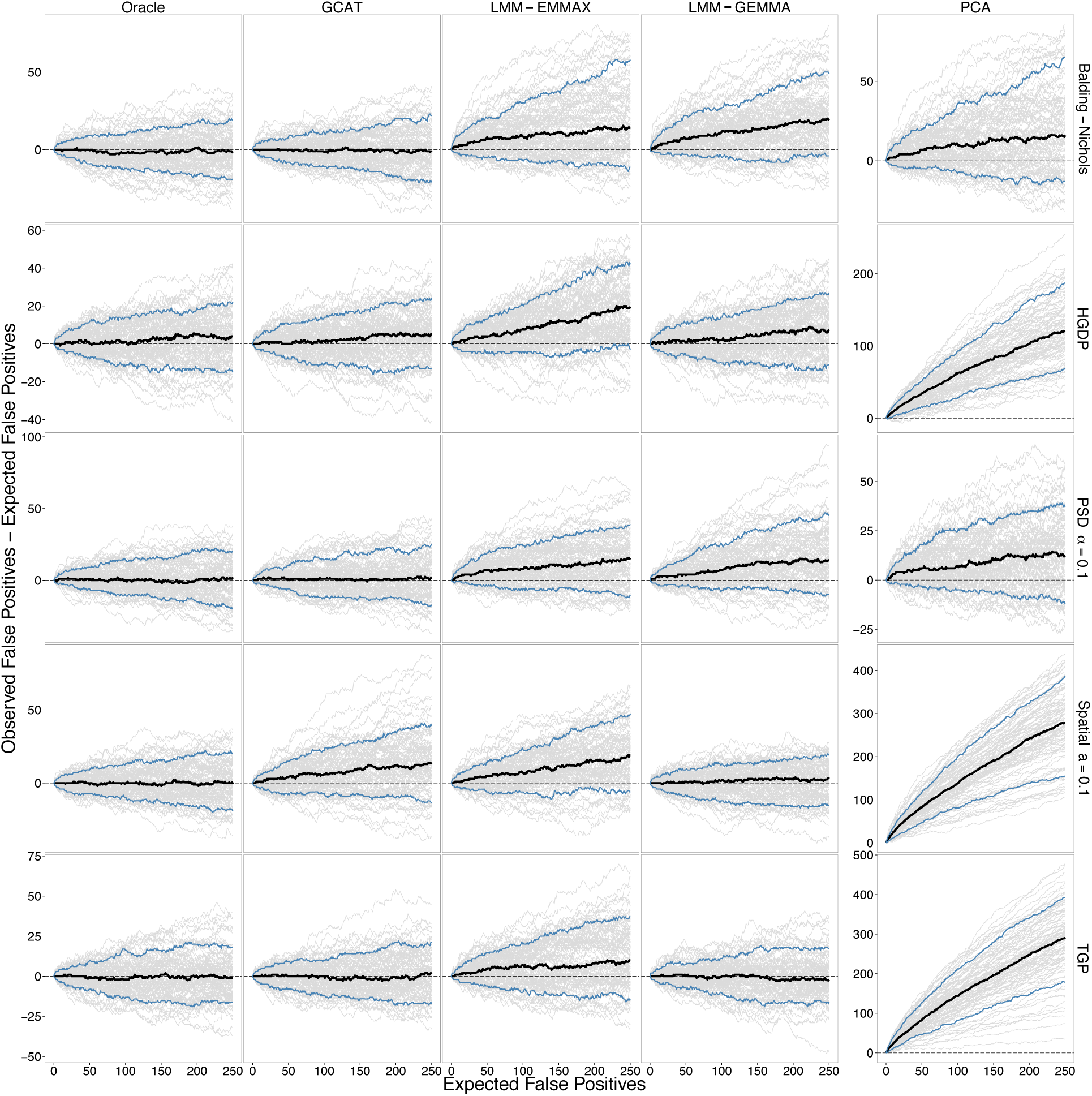
Performance of association testing methods. One-hundred quantitative trait GWAS studies were simulated in each of the Balding-Nichols, HGDP, TGP, PSD (*α* = 0.1), and Spatial (*a* = 0.1) simulation scenarios (see Online Methods for definitions of each) to compare the Oracle, GCAT (proposed), LMM-EMMAX, LMM-GEMMA, and PCA testing methods. The variance contributions to the trait are genetic=5%, non-genetic=5%, and noise=90%. The difference between the observed number of false positives and expected number of false positives is plotted against the expected number of false positives under the null hypothesis of no association for each simulated study (grey lines), the average of those differences (black line), and the middle 90% (blue lines). All simulations involved *m* = 100,000 SNPs, so the range of the x-axis corresponds to choosing a significance threshold of up to p-value ≤ 0.0025. The difference on the y-axis is the number of “spurious associations.” PCA is shown on a separate y-axis since it usually has a much larger maximum than the other methods. The Oracle method is where the true population structure parameters are inputted into the proposed test (see Results), which we have theoretically proven always corrects for structure (see Supplementary Note).

We found from using the distributed binary executable EMMAX software and our own implementation that EMMAX required a 10-fold increase in computational time over the proposed method and PCA when analyzing *n* = 5000 individuals. Therefore, it was not reasonable to apply EMMAX to all 3300 simulated GWAS data sets. We limited comparisons with EMMAX to five representative structure configurations. GEMMA was computationally more efficient, though still significantly slower than GCAT or our implementation of PCA.

Supplementary Figs. 1-8 show results from the remaining set of simulations from all 33 simulation configurations. Due to the computational constraints mentioned above for EMMAX, the additional simulations feature only results from GEMMA for LMM methods.

In comparing the statistical power among the methods (Supplementary Figs. 9-17), we found that the Oracle, GCAT, and PCA performed similarly well, while the two LMM methods sometimes showed a loss or gain in power depending on the scenario. We also carried out analogous simulations on binary traits simulated (from trait model equation (2) in Online Methods) and we found that all methods performed similarly well in terms of producing correct p-values that were robust to structure. This result agrees with the comparisons made between PCA and a linear mixed effects model in Astle and Balding (2009) [5].

### Analysis of the Northern Finland Birth Cohort Data

We applied the proposed method to the Northern Finland Birth Cohort (NFBC) genome-wide association study data [12], which includes several metabolic traits and height (Supplementary Figure 18). This study has also been analyzed by the LMM and PCA methods, as well as a standard analysis uncorrected for structure [10]. We carried out association analyses with the proposed method on the 10 traits that were also analyzed using the other methods (Table 1). After processing the data, including filtering for missing data, minor allele frequencies, and departures from Hardy-Weinberg equilibrium, the data were composed of *m* = 324,160 SNPs and *n* = 5027 individuals (Supplementary Note). The LFA model of population structure was estimated from a subset of the data where markers were at least 200 kbp apart.

**Table 1.** Number of significant loci at genome-wide significance (p-value < 7.2×10^−8^) for each of the 10 traits from the Northern Finland Birth Cohort data. Each method was performed with a subsequent genomic control inflation factor correction applied (denoted by +GC). The counts for LMM+GC, PCA+GC, and Uncorr+GC were obtained from Table 2 in Kang et al. (2010). In this case LMM is EMMAX-LMM.

| Trait | Abbreviation | GCAT+GC | LMM+GC | PCA+GC | Uncorr+GC |
| --- | --- | --- | --- | --- | --- |
| Body Mass Index | BMI | 0 | 0 | 0 | 0 |
| C-reactive Protein | CRP | 2<sup>†</sup> | 2 | 2 | 2 |
| Diastolic blood pressure | DBP | 0 | 0 | 0 | 0 |
| Glucose | GLU | 3 | 2 | 2 | 2 |
| HDL Cholesterol | HDL | 4 | 4 | 2 | 4 |
| Height | Height | 1 | 0 | 0 | 0 |
| Insulin | INS | 0 | 0 | 0 | 0 |
| LDL Cholesterol | LDL | 4 | 3 | 3 | 3 |
| Systolic blood pressure | SBP | 0 | 0 | 0 | 0 |
| Triglycerides | TG | 2 | 3 | 2 | 2 |
|  | Total | 16 | 14 | 11 | 13 |

<sup>†</sup>Result when the Box-Cox transformation was not applied to the CRP trait. The result is 1 when the transformation is applied.

Most traits showed only approximate Normal distributions, so we applied a Box-Cox Normal transformation to all traits so that they satisfy the model assumptions. We noted that C-reactive Protein (CRP) and Triglycerides (TG) traits followed an exponential distribution more closely, so it was unnecessary to transform these two traits. The developed theory can be extended to exponential distributed quantitative traits as well.

The 20 most significant SNPs for each of the 10 traits are shown in Supplementary Table 1. Kang et al. (2010) utilized a genome-wide significance threshold of p-value < 7.2×l0^−8^ as proposed in ref. [16], so we also utilized this threshold for comparative purposes. The numbers of loci found to be significant for each method are shown in Table 1. Whereas our proposed method identifies 16 significant loci, the other methods identify 11 to 14 loci.

We identified three new loci that were not identified by the other methods. None of the other methods identified any significant associations for the height trait. However, we identified rs2814982 on chromosome 6 as being statistically associated with height (Supplementary Table 1). This SNP is located ~70kbp from another SNP, rs2814993, which has been associated with skeletal frame size in a previous study [13]. Additionally, rs2814993 was the fifth most significant SNP for height. For the LDL cholesterol trait, we identified a significant association with rs11668477, which was significantly associated with LDL cholesterol in a different study [17]. Finally, there were significant associations between the glucose (GLU) trait and a cluster of SNPs (rs3847554, rs1387153, rs1447352, rs7121092) proximal to the MTNR1B locus; variation at this locus has been associated with glucose in a previous study [18].

As described in Sabatti et al. (2009) [12], the NFBC data show modest levels of inflation due to population structure as measured by the genomic control inflation factor (GCIF) [19] of test statistics from an uncorrected analysis. The population structure present among these individuals may be subtler and manifested on a finer scale than other settings. Noting that the GCAT approach does not attempt to adjust for a polygenic background, the GCIF values calculated for the proposed method (Supplementary Table 2) were found to be in line with what is expected for polygenic traits where no structure is present [20], providing evidence that the proposed method adequately accounts for structure.

## DISCUSSION

We considered models of quantitative and binary traits involving genetic effects and non-genetic effects in the presence of arbitrarily complex population structure. We allowed for the nongenetic effects to be confounded with population genetic structure since structure, ancestry, geography, lifestyle, and environment – all factors potentially involved in complex traits – may be highly dependent with one another. A mathematical argument showed that under these models, it is most reasonable to account for this confounding in the genotypes, but it is not tractable to do so in the non-genetic effects. This follows because we have many instances of genotypes that can be jointly modeled to provide reliable estimates of structure, but the nongenetic effects are never directly observed and we do not have repeated instances of them. In general it is not possible to estimate a latent variable that accounts for the confounding between structure and non-genetic effects.

These observations led us to propose an inverse regression approach to testing for associations, where the association is tested by modeling genotype variation in terms of the trait plus model terms accounting for structure. In this model, the terms accounting for structure were based on the logistic factor analysis (LFA) approach that we have proposed [14], although the general form of the association test can incorporate other methods that estimate population structure. We mathematically proved under general assumptions that the trait term in the model is non-zero only when the genetic marker is truly associated with the trait, regardless of the population structure. We demonstrated that the implemented test properly accounts for structure in a large body of simulated studies that included a wide range of population structures. We also applied the method to 10 traits from the Northern Finland Birth Cohort genome-wide association study. The proposed method identified three new loci associated with the traits, including being the only method among those we considered that identifies a locus associated with the height trait. Overall, we showed that the proposed method compares favorably to existing methods and we also noted that it has favorable computational requirements compared to existing methods.

As GWAS increase in sample size and levels of complexity of population structure, it is important to develop methods that properly account for structure and that scale well with sample size. Whereas we found that the popular principal components adjustment does not properly account for structure, we also found that the mixed model approach performs reasonably well. However, the mixed model approach involves estimating a *n*×*n* kinship matrix (where *n* is the number of individuals in the study) and its current implementation does not scale well with sample size. The kinship matrix quickly becomes computationally unwieldy when *n* grows large, and the possibility of the estimated kinship matrix becoming overwhelmed by noise is a concern [21]. In the Northern Finland Birth Cohort data, the mixed model approach required us to estimate 12 million parameters, whereas the proposed method involved estimating 25-thousand parameters, a ~500-fold decrease. A study involving *n* = 10,000 individuals with the same complexity of structure requires estimating about 50-million parameters in the mixed model kinship matrix, whereas the proposed method requires estimating 50-thousand parameters, a ~1000-fold decrease. In addition, estimating the structure in the proposed method primarily uses singular value decomposition, for which a rich literature of computational techniques exists. We utilized a Lanczos bidiagonalization algorithm [22], which scales approximately linearly with respect to *n* for *d* ≪ *n*, where *d* is the number of latent variables used in the LFA model of population structure (Online Methods). The proposed method is well equipped to scale to massive GWAS and can take advantage of future advances for computing singular value decomposition.

The key assumption to verify in utilizing the proposed GCAT approach is that population structure observed in the SNP genotypes is adequately modeled and estimated. One can test for associations among SNPs that show convincing empirical evidence that the model of structure is reasonably well-behaved; this can be directly tested on the genotype data as previously demonstrated in our logistic factor analysis (LFA) model of structure [14]. For example, on the Northern Finland Birth Cohort Study, we empirically verified that utilizing the LFA model with dimension *d* = 6 accounted for structure reasonably well for the great majority of SNPs. The linear mixed effects model (LMM) approach and principal components (PCA) approach make trait model assumptions that may be difficult to verify in practice (Online Methods and Supplementary Note). In cases where the probabilistic model that we assume is not validated on the data, a different model should be utilized. For example, our probabilistic model does not account for closely related individuals.

We anticipate that the proposed genotype conditional association test (GCAT) will be useful for future studies. The framework we have developed should facilitate its extension to traits modeled according to distributions not considered here while maintaining our theoretical proof that the test accounts for population structure in the presence of non-genetic effects also confounded with structure.

## URLs

The proposed method has been implemented in open source software, available at http://github.com/StoreyLab/gcat/.

## Acknowledgments

This research was supported in part by NIH grant R01 HG006448. The Northern Finland Birth Cohort data were collected by the STAMPEED: Cardiovascular Health Study (CHS) GWAS, made available through dbGaP Study Accession phs000226.v2.p1. A full list of contributors to the STAMPEED study can be found at its dbGaP web site.

## Author Contributions

JDS designed the study and wrote the manuscript. MS and JDS developed statistical theory and methods. WH developed statistical algorithms, simulated and analyzed data, and implemented software.

## METHODS

### Population Structure Model

Suppose that there are *n* individuals, each with *m* measured SNP genotypes. The genotype for SNP *i* in individual *j* is denoted by *x_ij_* ∈ {0,1,2}, *i* = 1,2, …, *m*, *j* = 1,2, …, *n*. We collected these SNP genotypes into an *m*×*n* matrix ***X***, where the (*i*, *j*) entry is ***x****_ij_*. We denote the genotypes for individual *j* by ***x^j^*** = (*x*_1*j*_, *x*_2*j*_, …, *x_mj_*)*^T^*.

We utilize our recently developed framework that flexibly models complex population structures for diallelic loci [14]. As described in Results, ***z*** is an unobserved latent variable that is assumed to capture heterogeneity in allele frequencies among individuals and can be interpreted as capturing the effect of population structure. (Note that, as described in Results, ***z*** also captures information on non-genetic contributions to the trait, such as those related to lifestyle and environment.) For a SNP *i*, the allele frequency *π_i_* can be viewed as being a function of ***z***, *π_i_*(***z***). For a random sample of *n* individuals, we therefore have implicitly sampled unobserved ***z***_1_, ***z***_2_, …, ***z****_n_* with resulting allele frequencies *π_i_*(***z***_1_), *π_i_*(***z***_2_), …, *π_i_*(***z****_n_*) for SNP *i*. In Hao et al. (2013) [14], we formulate and estimate a model for *m* SNPs simultaneously while providing a flexible parameterization of the form of *π_i_*(***z***).

For shorthand, *π_ij_* ≡ *π_i_*(***z****_j_*) is the allele frequency for SNP *i* conditioned on the ancestry state of individual *j*. The *π_ij_* values may be called “individual-specific allele frequencies” [14]. These allele frequencies can be collected into an *m*×*n* matrix ***F***, where the (*i*, *j*) entry is *π_ij_*. Note that E[*x_ij_*/2|***z****_j_*] = *π_ij_*, and when Hardy-Weinberg equilibrium holds, *x_ij_*|***z****_j_* ~ Binomial(2, *π_ij_*). We utilize the framework from Hao et al. (2013) [14], called “logistic factor analysis” (LFA), that allows the simultaneous estimation of all *π_ij_* from a given genotype data set ***X***. Specifically, it provides estimates of latent variables that form a linear basis of the 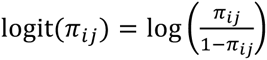 quantities, which turns out is the most convenient scale on which to estimate a model of structure for the proposed testing framework. It should be noted that other well-behaved estimates of *π_ij_* may be utilized as well. Further details are provided in Supplementary Note.

### Trait Models

We assume a trait (either quantitative or binary) has been measured on each individual, which we denote by *y_j_*, *j* = 1,2, …, *n*. We consider the following models of quantitative and binary traits. We write the trait models in terms of additive genetic effects, but the framework can be extended to account for dominance models and interactions, and the models can also incorporate adjustment variables that capture known sources of trait variation.

The quantitative trait model is

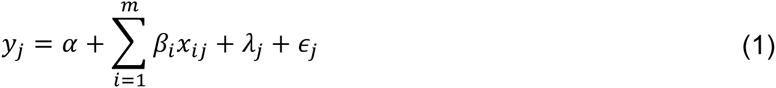

where *β_i_* is the genetic effect of SNP *i* on the trait, *λ_j_* is the random non-genetic effect, and *ϵ*_*j*_ is the random noise variation. To allow the interdependence of structure, lifestyle, and environment, we assume that ***x****^j^* = (*x*_l*j*_, …, *x_m,j_*)*^T^*, *λ_j_*, and 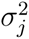 may all be functions of ***z****_j_*. We assume that 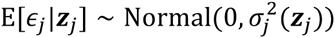, which allows for heteroskedasticity of the random noise variation. The distribution of *λ_j_* can remain unspecified, although we assume that *λ_j_* and ***z****_j_* may be dependent random variables. The population genetic model summarized shows how the distribution of 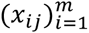, depends on ***z****_j_*. Without having observed ***z****_j_*, it follows that 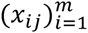, *λ_j_*, and *ϵ_j_* are dependent random variables; however, we assume that conditional on ***z****_j_*, these random variables are independent.

The binary trait model is

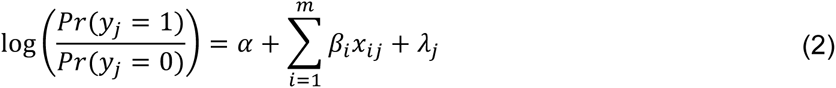

where again *β_i_* is the genetic effect of SNP *i* on the trait, *λ_j_* is the non-genetic effect, and we allow for the case that ***x***^*j*^ and *λ_j_* may be dependent due to the common confounding latent variable ***z****_j_* as described for the quantitative trait model.

We have shown that the linear mixed effects model and principal components approaches involve more restrictive assumptions about the trait models utilized in testing for associations (Supplementary Note).

### Association Test Immune to Population Structure

We have derived a statistical hypothesis test of association that is equivalent to testing whether *β_i_* = 0 for each SNP *i* in the above trait models (1) and (2), and whose null distribution does not depend on structure or the non-genetic effects correlated with structure, making it immune to spurious associations due to structure (Supplementary Note). Specifically, the test allows for general levels of complexity in structure because the test is based on adjusting for structure according to individual-specific allele frequencies.

We have proved a theorem (Supplementary Note) that shows that *β_i_* = 0 in models (1) and (2) implies that *b_i_* = 0 in the following model:

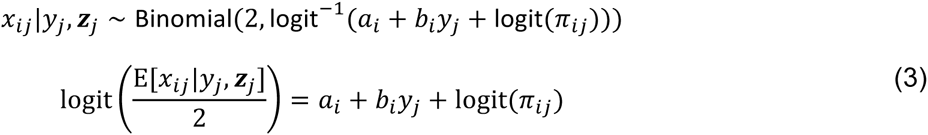

for all *j* = 1,2, …, *n*. This establishes a model that can be used to test for associations in place of models (1) and (2). Note that the non-genetic effects, heteroskedasticity, and polygenic background do not appear in the above model used to test for associations. This is important because under our general assumptions, these terms can be difficult or even impossible to estimate in practice. Furthermore, testing for association under this model means that the test will have a valid null distribution regardless of the form of the non-genetic effects, heteroskedasticity, and polygenic background.

As fully detailed in Supplementary Note, an association statistic whose null distribution is known can be constructed by testing whether *b_i_* = 0 in the above model, which we have shown is a valid test if *β_i_* = 0 in traits models (1) and (2). Briefly, the testing procedure works as follows:

1. Formulate and estimate a model of population structure that provides well-behaved estimates of the logit(*π_ij_*) values. We specifically use the logistic factor analysis (LFA) approach of ref. [14], which has been shown to provide a accurate linear basis of the logit(*π_ij_*) values.
2. For each SNP *i*, perform a logistic regression of the SNP genotypes on the trait values plus the model terms that estimate the 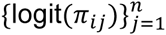 values. Also, perform a logistic regression of the SNP genotypes on only the model terms that estimate 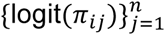, where the trait is now excluded from the fit. These two model fits are compared via a likelihood ratio statistic, where the larger the statistic, the more evidence there is that *b_i_* ≠ 0.
3. Calculate a p-value for each SNP, which is done based on our result that when the null hypothesis of no association is true, *β_i_* = 0 in models (1) and (2), then the above statistic follows a 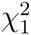 distribution for large sample sizes.

In our implementation, *d* estimated logistic factors (from LFA [14]) are included as covariates, which serve as the model terms that estimate the 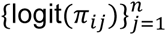 values.

We call our proposed test the “genotype-conditional association test” (GCAT). As a general concept, such an approach is sometimes called an inverse regression model because the trait and genotype are reversed in the regression.

### Simulated Data

The complete simulation study on quantitative traits involved population structure constructed in 11 different ways for each of three different apportionments of variance among genetic effects, non-genetic effects, and random variation that all contribute to variation in the trait. Therefore, each configuration involved a constructed allele frequency matrix ***F*** and values assigned to variances 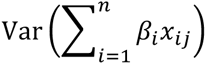, 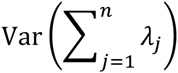, and Var(*ϵ_j_*) from model (1). For each of these 33 = 11×3 configurations, we simulated 100 GWAS data sets, for a grand total of 3300 studies.

We simulated allele frequencies: (i) from the Balding-Nichols model [23] based on allele-frequency and F_st_ estimates calculated on the HapMap data set (*Balding-Nichols*); (ii) subject to structure estimated from two real data sets: the Human Genome Diversity Project (*HGDP*) and the 1000 Genomes Project (*TGP*); (iii) at four different levels of admixture by varying the parameter α (defined in Supplementary Note) in the Pritchard-Stephens-Donnelly (*PSD*) model [24], which is an extension of the Balding-Nichols model; and (iv) for four different types of spatially defined structure (*Spatial*) by varying the parameter *a* (defined in Supplementary Note). We intentionally simulated challenging population structures, having in mind that future GWAS such as the forthcoming “Genotype Tissue Expression” program (GTEx) data may involve particularly challenging forms of structure.

In order to provide an extra challenge to the proposed test, we simulated the allele frequencies from a model that differs from the LFA model (equation 4 in Supplementary Note). We generated allele frequencies parameterized by ***F*** = **Γ*S***, where ***F*** is the matrix of *π_ij_* values, **Γ** is an *m*×*d* matrix and ***S*** is the *d*×*n* matrix that encapsulates the structure (with *d* = 3). This model captures as special cases the Balding-Nichols model and the PSD model [14]. It was also intended to provide an advantage to the PCA and LMM methods because the structure is manifested on the observed genotype scale [14], which is the same scale on which both methods estimate structure.

We simulated 10 truly associated SNPs whose effect sizes are distributed according to a Normal distribution. All genotypes were simulated to be in linkage equilibrium so that true and false positives are unambiguous. We set the variances 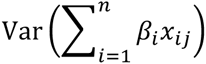, 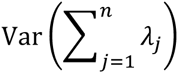, and Var(*ϵ_j_*) to be: (5%, 5%, 90%), (10%, 0%, 90%), and (10%, 20%, 70%). Setting these variances enforced a certain overall level of genetic contribution to the trait; therefore our simulation study results were minimally affected by the choice of 10 truly associated SNPs and the Normal distribution on their effect sizes. In each simulation scenario, we simulated data for *m* = 100,000 SNPs and *n* = 5000 individuals, except HGDP necessarily restricted us to *n* = 940 individuals and TGP to *n* = 1500 individuals. The dimension of the structure was set to *d* = 3, although we carried out the same simulations for *d* = 6 and the results were quantitatively very similar and qualitatively equivalent.

Additional details on the simulations can be found in Supplementary Note.

## Supplementary Materials

### Supplementary Note

This supplementary note assumes the reader is familiar with the details and mathematical notation from Online Methods.

#### 1 Logistic Factor Analysis (LFA)

When forming a latent variable model of structure, where the goal is to make minimal assumptions about the underlying structure, there are benefits to modeling logit(*π_ij_*) in terms of a latent variable model instead of *π_ij_* directly [1]. The quantity logit(*π_ij_*) = log(*π_ij_/*(1 − *π_ij_*)) is called the “natural parameter” of the distribution of *x_ij_* when we assume Hardy-Weinberg equilibrium so that *x_ij_* ~ Binomial(2, *π_ij_*). The quantity logit(*π_ij_*) occurs as a linear term in the log-likelihood of the data, and it is the target parameter in logistic regression because of its straightforward mathematical properties. This viewpoint also facilitates calculating the distribution of *x_ij_* given the structure, which is the essential challenge in accounting for structure in the proposed association testing framework.

In the association testing framework we have developed, it turns out that developing a latent variable model and estimate of the logit(*π_ij_*) is particularly appropriate. The approach is called “logistic factor analysis” (LFA). Let **L** be an *m* × *n* matrix with (*i, j*) element equal to logit(*π_ij_*). Consider the following parameterization:

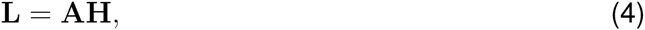

where **A** is an *m*×*d* matrix, **H** is a *d*×*n* matrix, and *d* ≪ *n*. The columns of **H** are independent, and column *j* captures the structure information for individual *j*. That is, Pr(*x_ij_*|**h***^j^*, ***z****_j_*) = Pr(*x_ij_*|**h***^j^*) where **h***^j^* is column *j* of **H**. Row *i* of **A** determines how SNP *i* is affected by structure. We have shown in ref. [1] that this model performs well in estimating structure resulting from discrete subpopulations, admixed populations, the Balding-Nichols model [2], the Pritchard-Stephens-Donnelly model [3], and models of spatially oriented structure.

In practice, **H** will be unknown, so it must be estimated. We have developed a method called logistic factor analysis (LFA) that we have shown to estimate **H** well [1]. Specifically, the LFA estimate 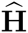 has been shown to span the same space as the true **H** at a high level of accuracy, which implies that replacing **H** with 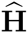 in the above equations yields nearly identical results. The accuracy of 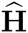 in estimating **H** has been demonstrated even when the individual-specific allele frequencies are not directly constructed from model (4), **L** = **AH**.

#### 2 Proposed Association Testing Framework

We have derived a statistical hypothesis test of association that is equivalent to testing whether *β_i_* = 0 for each SNP *i* in the trait models (1) and (2) (defined in Online Methods), and whose null distribution does not depend on structure or the non-genetic effects correlated with structure, making it immune to spurious associations due to structure. Specifically, the test allows for general levels of complexity in structure because the test is based on adjusting for structure according to individual-specific allele frequencies.

##### A Model of Genetic Variation Given the Trait and Structure

As a first step, we have proved a theorem (see below) that shows that *β_i_* = 0 in models (1) and (2) implies that *b_i_* = 0 in the following model:

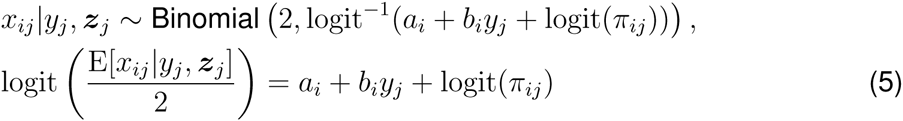

for all *j* = 1, 2, …, *n*. This establishes a model that can be used to test for associations in place of models (1) and (2).

There are a few important details to note. First, the variables *λ_j_*, 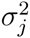, and (*x_kj_*)*_k_*_≠*i*_ do not appear in the model. This is important because it is impossible to estimate *λ_j_* and 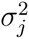 in the typical setting, and we will also typically not know the polygenic 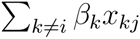 component of the model. Second, the genotype variation is being modeled in terms of the trait variation, instead of the other way around. It is initially counter-intuitive because almost all association tests involve modeling the trait in terms of the SNP genotypes. As explained in more detail below, this reversal is crucial for adjusting the probability distribution of *x_ij_* according to structure, and for eliminating the need to estimate *λ_j_*, 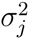, and (*β_k_*)*_k_*_≠*i*_.

We call our proposed test the “genotype conditional association test” (GCAT). The model we propose to utilize is sometimes called an inverse regression model because we utilize E[*x*|*y*] rather than E[*y*|*x*].

##### Proposed Test Conditional on Individual-Specific Allele Frequencies

As a second step, we have derived a test-statistic to test whether *b_i_* = 0 in model (3) (defined in Online Methods) whose null distribution is immune to structure. The log-likelihood function of the parameters given individual *j* is

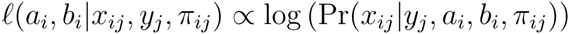

where the probability on the right-hand-side is calculated according to model (3). The log-likelihood of all *n* individuals is

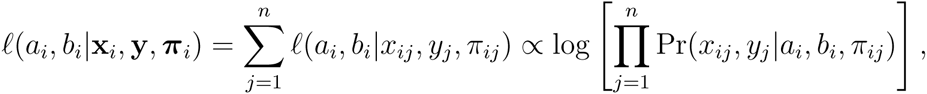

where *π_i_* = (*π_i_*_1_, *π_i_*_2_, …, *π_i,n_*) and **y** = (*y*_1_*, y*_2_, …, *y_n_*). The test statistic we utilize is a generalized likelihood ratio test statistic [4]:

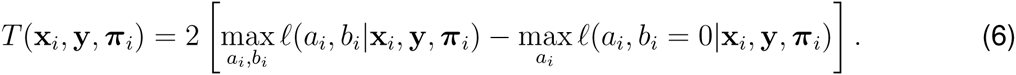

The log-likelihood is maximized by performing a logistic regression of all *n* observed genotypes for SNP *i* on the right hand side of model (3). We have proven a theorem below that shows that when *β_i_* = 0 in models (1) or (2), the null distribution of this test statistic is 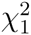, regardless of the values of *π_ij_*, (*x_kj_*)*_k_*_≠*i*_, (*β_kj_*)*_k_*_≠*i*_, *λ_j_*, and 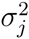 for *j* = 1, 2, …, *n* in models (1) and (2).

##### Proposed Test In Terms of LFA Model

As a third step, we have extended the above results to the case where the individual-specific allele frequencies are unknown and must be estimated. This requires a model of the individual-specific allele frequencies, and we utilize model (4) so that 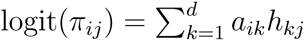. First, assume that **H** from model (4) is known. We have proved that *β_i_* = 0 in models (1) and (2) implies *b_i_* = 0 in the following model:

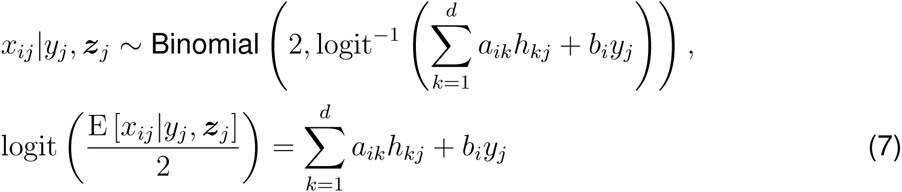

for all *j* = 1, 2, …, *n*, where **h***^j^* is column *j* of **H** and it is noted that without loss of generality we let *h_dj_* = 1 making *a_id_* an intercept term. The test-statistic used to test for an association between SNP *i* and the trait is the following generalized likelihood ratio test statistic:

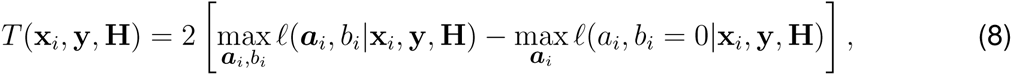

where ***a****_i_* = (*a_i_*_1_, *a_i_*_2_, *…, a_i,d_*). The log-likelihoods in this test statistic are maximized by performing a logistic regression of all *n* observed genotypes for SNP *i* on the right hand side of model (7) on all *n* individuals. As the previous case, we have proven a theorem below that shows that when *β_i_* = 0 in models (1) or (2), the null distribution of this test statistic is 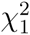, regardless of the values of *π_i_*, (*x_kj_*)*_k_*_≠*i*_, *β*_−*i*_, *λ*, and *σ*^2^ in models (1) and (2).

The proposed test utilizes LFA to form an estimate 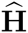, replaces **H** with 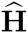, and carries out the test using model (7) and test statistic (8): 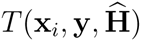. This approach directly allows the simultaneous estimation of ***a****_i_* and *b_i_* for each SNP *i* under the unconstrained model and the estimation of ***a****_i_* with *b_i_* = 0 under the constraints of the null hypothesis. Because of this, the test allows the uncertainty of the *m* × *d* unknown parameters of **A** to be taken into account and it allows *b_i_* to be competitively fit with ***a****_i_* under the unconstrained, alternative hypothesis model.

Another approach is to first carry out estimation of **F** by whatever method the analyst finds appropriate and then base the test on statistic (6) with the *π_ij_* replaced with the estimates 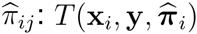. This has the advantage that it allows for a much broader class of methods to estimate **F**, but it may be more conservative than the above implementation because *b_i_* is not competitively fit with the *π_ij_* under the unconstrained model. In this case, **F** may be estimated in a manner that allows for fine-scale levels of inter-individual coancestry and locus-specific models of structure without relying on the lower *d*-dimensional factorized model **L** = **AH** that we used here.

##### Proposed Test Under the Alternative Hypothesis

The proposed association test is based on models (3) and (7). Even though we have proved that the test is immune to population structure, it is also important to demonstrate that the test has favorable statistical power to identify true associations. We have shown that the 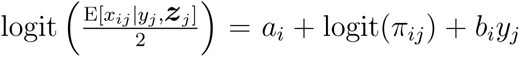 is a tractable approximation of the model under general configurations of a true alternative hypothesis for SNP *i* where *β_i_* ≠ 0 (see below). This provides the beginnings of a mathematical framework for characterizing the power of the test.

#### 3 Theorems and Proofs

Because *x_ij_*|***z****_j_* ~ Binomial(2*, π_i_*(***z****_j_*)) where we write *π_ij_ ≡ π_i_*(***z****_j_*), it follows that Pr(*x_ij_*|*π_ij_*, ***z****_j_*) = Pr(*x_ij_*|*π_ij_*). We assume that Pr(*x_ij_*|**h***^j^*, ***z****_j_*) = Pr(*x_ij_*|**h***^j^*); in other words, all information about the influence of population structure on the genotypes of individual *j* is captured through column *j* of **H**. It therefore follows that Pr(*x_ij_*|*π_ij_*, **h***^j^*, ***z****_j_*) = Pr(*x_ij_*|*π_ij_*, **h***^j^*) = Pr(*x_ij_*|*π_ij_*). We also assume that the SNP genotypes are mutually independent given the structure (which also implies the set of SNPs we consider are in linkage equilibrium, given the structure). These assumptions yield the following equalities:

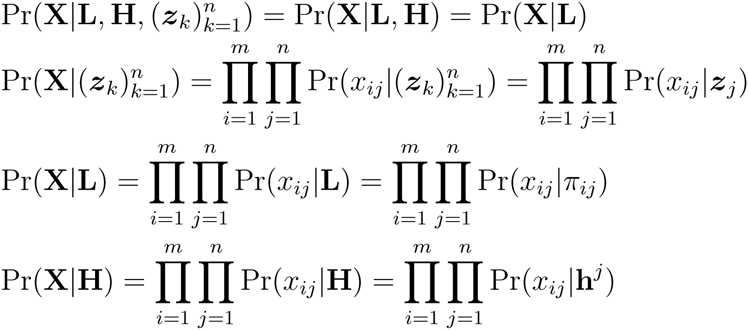

##### Theorem 1

*Suppose that y_j_ is distributed according to model* (1) *or* (2)*, x_ij_*|*π_ij_* ~ *Binomial*(2*, π_ij_*) *as parameterized above, and the SNP genotypes are mutually independent given the structure as detailed above. Then β_i_* = 0 *in models* (1) *or* (2) *implies that b_i_* = 0 *in model* (3).

*Note:* We provide two proofs of this theorem because both provide relevant insights. The first version gives insight into the probabilistic mechanism underlying the proposed approach and has some generality beyond the modeling assumptions made here. The second version directly shows how the terms in models (1) and (2) relate to those in model (3).

**Proof (version 1):** When *β_i_* = 0, it follows that Pr(*y_j_*|(*x_kj_*)*_k_*_≠*i*_, *x_ij_*, ***z****_j_*) = Pr(*y_j_*|(*x_kj_*)*_k_*_≠*i*_, ***z****_j_*) by the assumptions of models (1) and (2). Noting that Pr((*x_kj_*)*_k_*_≠*i*_|*x_ij_*, ***z****_j_*) = Pr((*x_kj_*)*_k_*_≠*i*_|***z****_j_*) by the conditional independence assumption, we have:

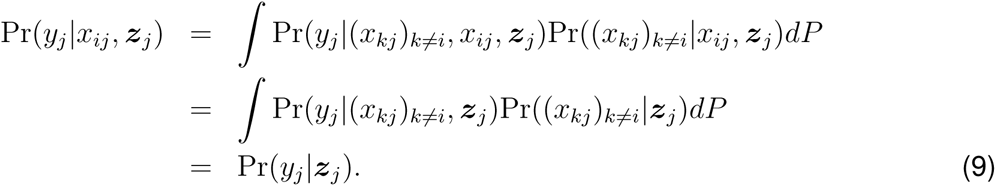

By Bayes theorem we have

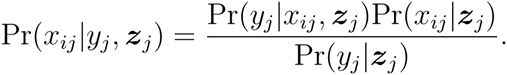

Since Pr(*y_j_*|*x_ij_*, ***z****_j_*) = Pr(*y_j_*|***z****_j_*), this implies that Pr(*x_ij_*|*y_j_*, ***z****_j_*) = Pr(*x_ij_*|***z****_j_*) and it follows that *b_i_* = 0 in model (3).

**Proof (version 2):** For either model (1) or (2), it follows that

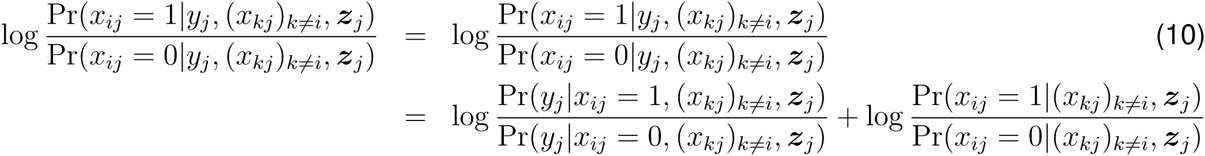

and similarly

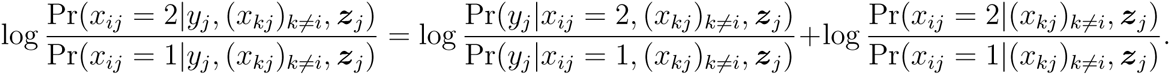

By the assumptions detailed above, we have Pr(*x_ij_*|(*x_kj_*)*_k_*_≠*i*_, ***z****_j_*) = Pr(*x_ij_*|*π_ij_*) and therefore:

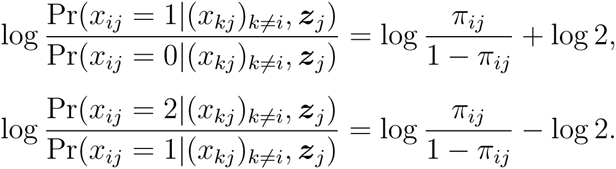

Under the *quantitative trait* model (1), it follows that

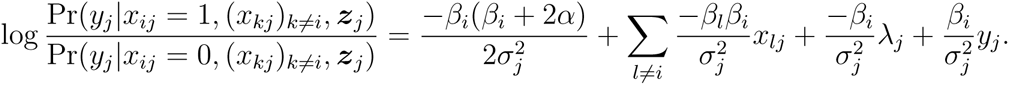

Plugging this back into equation (10) shows that

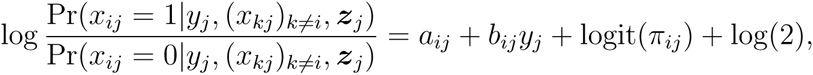

where 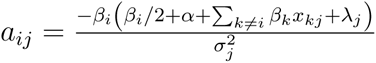 and 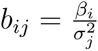. Following analogous steps, we find that

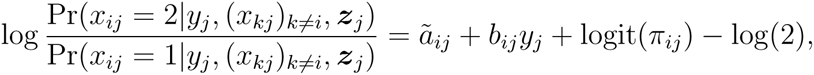

where 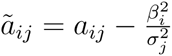. When *βi* = 0 in model (1), then 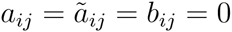.

Under the *binary trait* model (2), it follows that

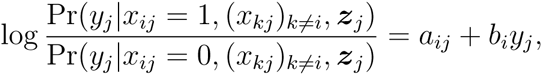

where 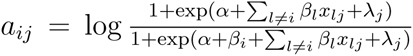 and *b_i_* = *β_i_*. Plugging this back into equation (10) shows that

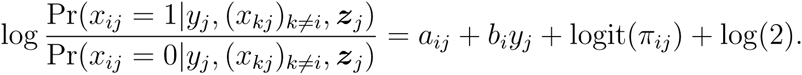

Following analogous steps, we find that

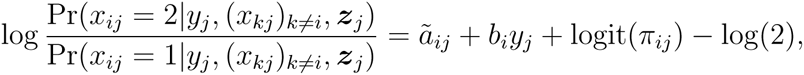

where 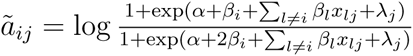. When *β_i_* = 0 in model (2), then 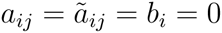.

Putting these together, we have that when *β_i_* = 0 in models (1) or (2), then model (3) holds with *b_i_* = 0.

##### Corollary 1

*Suppose that the assumptions of Theorem 1 hold and additionally* 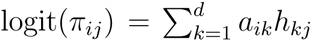. Then *β_i_* = 0 *in models* (1) *or* (2) *implies that b_i_* = 0 *in model* (7).

**Proof:** The proof is the same as that to Theorem 1, except we replace *π_ij_* with **h***^j^*.

##### Theorem 2

*Suppose that y_j_ is distributed according to model* (1) *or* (2) *and that x_ij_*|*π_ij_* ~ *Binomial*(2*, π_ij_*)*. If β_i_* = 0 *in models* (1) *or* (2)*, then the test-statistic T* (**x***_i_*, **y**, ***π****_i_*) *defined in* (6) *converges in distribution to* 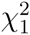 *as n* → ∞.

**Proof:** When *β_i_* = 0, then [*x_ij_*|*y_j_, π_ij_* ] ~ Binomial (2*, π_ij_*) by Theorem 1. It then follows that 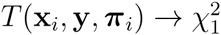 in distribution as *n* → ∞ by Wilks’ theorem [4].

##### Corollary 2

*Suppose that the assumptions of Theorem 1 hold and additionally* 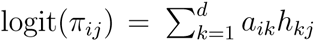. *If β_i_* = 0 *in models* (1) *or* (2)*, then the test-statistic T*(**x***_i_*, **y**, **H**) *defined in* (8) *converges in distribution to* 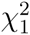 *as n* → ∞.

**Proof:** When *β_i_* = 0, then 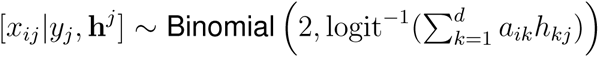 by Corollary 1. It then follows that 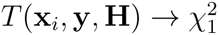 in distribution as *n* → ∞ by Wilks’ theorem [4].

#### 4 Proposed Model Under the Alternative Hypothesis

When the alternative model is true this means that *β_i_* ≠ 0. In this case it is worthwhile to characterize model (3) in terms of the distribution of *x_ij_*|*y_j_*, ***z****_j_*. Under trait models (1) or (2), it follows that:

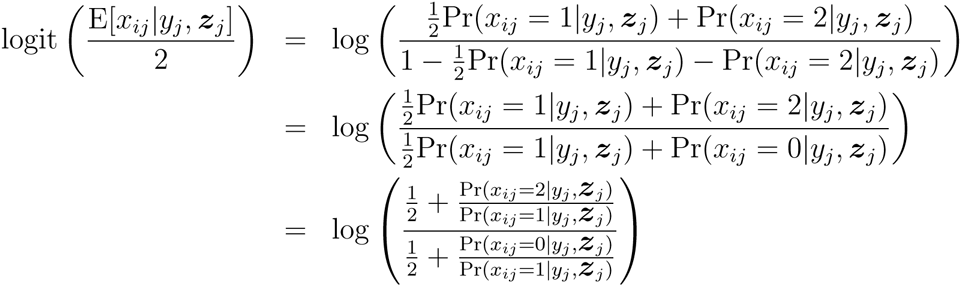

This implies that

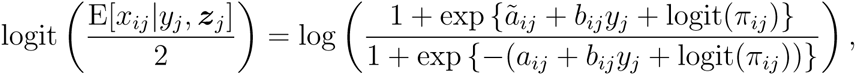

where under model (1) we have 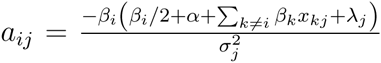, 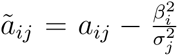, 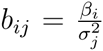 and under model (2) we have 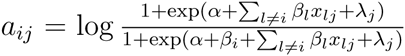, 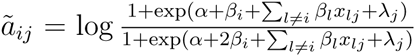, *b_ij_* = *β_i_*.

In the case that 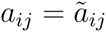, it is the case that

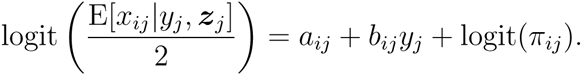

However, this exact equality is only the case when *β_i_* = 0. For the typical effect sizes seen in GWAS, it will nevertheless be true that 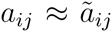, in which case the above functional form will be approximately true. This allows for an approximation that can be utilized in practice for power calcuations.

#### 5 Simulated Allele Frequencies

In order to simulate the *m* × *n* matrix of genotypes **X**, we first needed to simulate the *m* × *n* matrix of allele frequencies **F**. Recall that we model the allele frequencies by forming **L** = logit(**F**) and then utilizing the model **L** = **AH** from equation (4).

Instead of simulating allele frequencies from the **L** = **AH** model we use to perform the proposed association test, we instead simulated them from a different model to demonstrate the flexibility of the **L** = **AH** model. Specifically, we let **F** = **ΓS** where **Γ** is *m* × *d* and **S** is *d* × *n* with *d* ≤ *n*. The *d* × *n* matrix **S** encapsulates the genetic population structure for these individuals since **S** is not SNP-specific but is shared across SNPs. The *m* × *d* matrix **Γ** maps how the structure is manifested in the allele frequencies of each SNP. We have shown that the model **F** = **ΓS** includes as special cases discrete subpopulations, the Balding-Nichols model, and the Pritchard-Stephens-Donnelly model.

We formed **Γ** and **S** for the 11 different population structure configurations exactly as carried out in Hao et al. (2013) [1]. These constructions are summarized as follows from Hao et al. (2013).

##### Balding-Nichols Model (Balding-Nichols)

The HapMap data set was deliberately sampled to be from three discrete populations, which allowed us to populate each row *i* of Γ with three independent and identically distributed draws from the Balding-Nichols model: 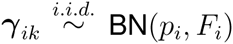, where *k* ∈ {1, 2, 3}. Each ***γ****_ik_* is interpreted to be the allele frequency for sub-population *k* at SNP *i*. The pairs (*p_i_, F_i_*) were computed by randomly selecting a SNP in the HapMap data set, calculating its observed allele frequency, and estimating its F_ST_ value using the Weir & Cockerham estimator [5]. The columns of **S** were populated with indicator vectors such that each individual was assigned to one of the three subpopulations. The subpopulation assignments were drawn independently with probabilities 60/210, 60/210, and 90/210, which reflect the subpopulation proportions in the HapMap data set. The dimensions of the simulated data were *m* = 100,000 SNPs and *n* = 5000 individuals.

##### 1000 Genomes Project (TGP)

We started with the TGP data set from Hao et al. (2013) [1]. The matrix **Γ** was generated by sampling 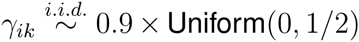 for *k* = 1, 2 and setting *γ_i_*_3_ = 0.05. In order to generate **S**, we computed the first two principal components of the TGP genotype matrix after mean centering each SNP. We then transformed each principal component to be between (0, 1) and set the first two rows of **S** to be the transformed principal components. The third row of **S** was set to 1, i.e. an intercept. The dimensions of the simulated data were *m* = 100,000 and *n* = 1500, where *n* was determined by the number of individuals in the TGP data set.

##### Human Genome Diversity Project (HGDP)

We started with the HGDP data set from Hao et al. (2013) [1] and applied the same simulation scheme as for the TGP scenario. The dimensions of the simulated data were *m* = 100,000 and *n* = 940, where *n* was determined by the number of individuals in the HGDP data set.

##### Pritchard-Stephens-Donnelly (PSD)

The PSD model assumes individuals to be an admixture of ancestral subpopulations. The rows of **Γ** were again created by three independent and identically distributed draws from the Balding-Nichols model: 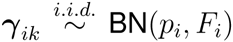, where *k* ∈ {1, 2, 3}. For this scenario, the pairs (*p_i_, F_i_*) were computed from analyzing the HGDP data set for observed allele frequency and estimated F_ST_ via the Weir & Cockerham estimate [5]. The estimator requires each individual to be assigned to a subpopulation, which were made according to the *K* = 5 subpopulations from the analysis in Rosenberg et al. (2002) [6]. The columns of **S** were sampled 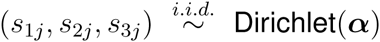 Dirichlet(***α***) for *j* = 1*, …, n*. There were four PSD scenarios with parameter values ***α*** = (0.01, 0.01, 0.01), ***α*** = (0.1, 0.1, 0.1), ***α*** = (0.5, 0.5, 0.5), and ***α*** = (1, 1, 1). ***α*** = (0.1, 0.1, 0.1) was chosen as the representative structure for Figure 2. The dimensions of the simulated data were *m* = 100,000 SNPs and *n* = 5000 individuals.

##### Spatial

We seek to simulate genotypes such that the population structure relates to the spatial position of the individuals. The matrix Γ was populated by sampling 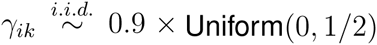 for *k* = 1, 2 and setting *γ_i_*_3_ = 0.05. The first two rows of **S** correspond to coordinates for each individual on the unit square and were set to be independent and identically distributed samples from Beta(*a, a*), while the third row of **S** was set to be 1, i.e. an intercept. There were four spatial scenarios with parameter values of *a* = 0.1, 0.25, 0.5, and 1. As *a* → 0, the individuals are placed closer to the corners of the unit square, while when *a* = 1, the individuals are distributed uniformly. *a* = 0.1 was chosen as the representative structure for Figure 2. The dimensions of the simulated data were *m* = 100,000 SNPs and *n* = 5000 individuals.

#### 6 Simulated Traits

For each of the 11 simulations scenarios, we generated 100 independent studies. For each study, **X** was formed by simulating *x_ij_* ~ Binomial(2*, π_ij_*) where **F** was constructed as described above. In order to simulate a quantitative trait, we needed to simulate *α*, 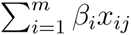, *λ_j_*, and *ϵ_j_* from model (1).

First, we set *α* = 0. Without loss of generality SNPs *i* = 1, 2, …, 10 were set to be true alternative SNPs (where *β_i_* ≠ 0); we simulated 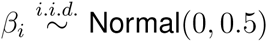 for *i* = 1, 2, …, 10. We set *β_i_* = 0 for *i >* 10. Note that **X** is influenced by the latent variables ***z***_1_, …, ***z****_n_* through **S** in the model **F** = **ΓS** described above. In order to simulate *λ_j_* and *ϵ_j_* so that they are also influenced by the latent variables ***z***_1_, …, ***z****_n_*, we performed the following:

1. Perform *K*-means clustering on the columns of **S** with *K* = 3 using Euclidean distance. This assigns each individual *j* to one of three mutually exclusive cluster sets 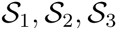 where 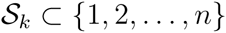.
2. Set *λ_j_* = *k* for all 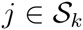 for each *k* = 1, 2, 3.
3. Let 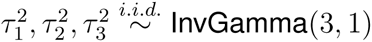 and set 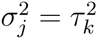 for all 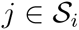 for each *k* = 1, 2, 3.
4. Draw 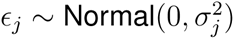 independently for *j* = 1, 2, …, *n*.

This strategy simulates non-genetic effects and random variation that manifest among *K* discrete groups over a more continuous population genetic structure defined by **S**. This is meant to emulate the fact that environment (specifically lifestyle) may partition among individuals in a manner distinct from, but highly related to population structure.

This yields three values 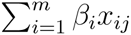, *λ_j_*, and *ϵ_j_* for each individual *j* = 1, 2, …, *n*. In order to set the variances of these three values to pre specified levels *ν*_gen_, *ν*_env_ and *ν*_noise_, we rescaled each quantity as follows:

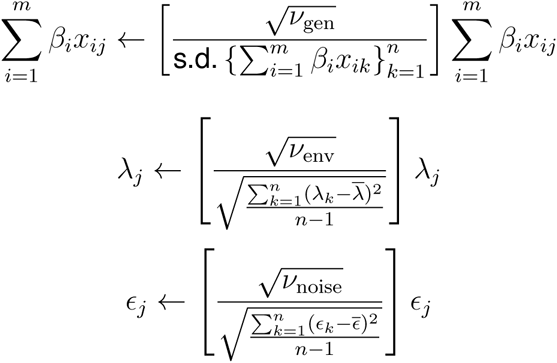

The trait for a given study was then formed according to

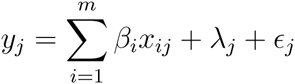

for *j* = 1, 2, …, *n*. For each of the 11 simulation scenarios, we considered the following three configurations of (*ν*_gen_*, ν*_env_*, ν*_noise_): (5%, 5%, 90%), (10%, 0%, 90%) and (10%, 20%, 70%).

In total, there were 11 different types of structures considered over three different configurations of genetic, environmental, and noise variances for a total of 33 settings. For each setting, we simulated 100 independent studies where each involved *m* = 100,000 SNPs and up to *n* = 5000 individuals.

#### 7 Northern Finland Birth Cohort Data

Genotype data was downloaded from dbGaP (Study Accession: phs000276.v1.p1). Individuals were filtered for completeness (maximum 1% missing genotypes) and pregnancy. (Pregnant women were excluded because we did not receive IRB approval for these individuals.) SNPs were first filtered for completeness (maximum 5% missing genotypes) and minor allele frequency (minimum 1% minor allele frequency), then tested for Hardy-Weinberg equilibrium 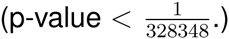. The final dimensions of the genotype matrix are *m* = 324, 160 SNPs and *n* = 5027 individuals.

A Box-Cox transform was applied to each trait, where the parameter was chosen such that the values in the median 95% value of the trait was as close to the normal distribution as possible. Indicators for sex, oral contraception, and fasting status were added as adjustment variables. For glucose, the individual with the minimum value was removed from the analysis as an extreme outlier. All analyses were performed with *d* = 6 logistic factors, which was determined based on the Hardy-Weinberg equilibrium method described in ref. [1]. The association tests were performed exactly as described in the main text.

#### 8 Linear Mixed Effects Model and Principal Component Analysis Approaches

In order to explain the assumptions made by the linear mixed effects model approach (LMM) and principal components approach (PCA), we first re-write model (1) as follows:

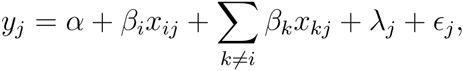

where the object of inference is *β_i_* for each SNP *i* = 1*, …, m*. As explained in Astle and Balding (2009) [7], these approaches assume that 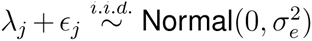, meaning that the non-genetic effects are independent from population structure and there is no heteroskedasticity among individuals.

The LMM approach also makes the assumption that we can approximate the genetic contribution by a multivariate Normal distribution:

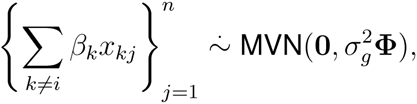

where **Φ** is the *n* × *n* kinship matrix. If we define 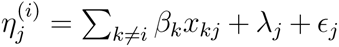, we can write the above model as

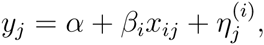

where it is assumed that 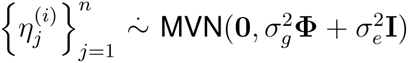. Since it is not the case in general that the 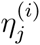 are identically distributed for all SNPs *i* = 1, …, *m*, one can either estimate a different pair of parameters 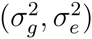 for each SNP or assume that these parameters change very little between SNPs. Since the former tends to be computationally demanding, algorithms such as EMMAX [8] propose to estimate a single pair of parameters 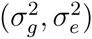 from a null model and then utilize this single estimate for every SNP. More recently, algorithms such as GEMMA have been proposed to relax this assumption [9].

The *n* × *n* kinship matrix **Φ** is estimated from the genotype data **X**. This involves the simultaneous estimation of (*n*^2^ − *n*)/2 parameters, which is particularly large for sample sizes considered in current GWAS (on the order of 10^8^ for *n* = 10,000). The uncertainty in the estimated **Φ** is typically not taken into account, and there is so far no regularization of the high-dimensional estimator of **Φ**. Unregularized estimates of large covariance matrices have been shown to be problematic [10, 11], a concern that is also applicable to estimates of **Φ**. Estimating 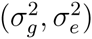 involves manipulations of the estimated **Φ** matrix, which can pose numerical challenges due to the fact that the estimated **Φ** is both high-dimensional and nonsingular. The LMM approach therefore makes assumptions that are important to verify for each given study and it involves some challenging calculations and estimations.

The PCA approach first calculates the top *d* principal components on a normalized version of the genotype matrix **X**. In the method proposed by Price et al. (2006) [12], these principal components are then regressed out of each SNP *i* and the trait (regardless of whether it is binary or quantitative). A correlation statistic is calculated between each adjusted SNP genotype and the adjusted trait, and the p-value that tests for equality to 0 is reported. As shown in Hao et al. (2013) [1], the top *d* principal components form a high-quality estimate of a linear basis of the allele frequencies *π_ij_*. Extracting the residuals after linearly regressing the genotype data for SNP *i* onto these principal components is equivalent to estimating the quantity *x_ij_* − *π_ij_*. Using the trait as the response variable in this regression adjustment is equivalent to estimating 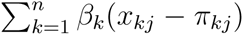 under the assumptions on the trait model given above (where this quantitative trait model is assumed regardless of whether the trait is quantitative or binary). Therefore, the association test carried out in the PCA approach implicitly involves an estimated form of the model:

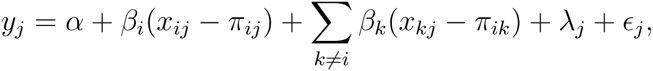

where it is assumed that *λ_j_* + *ϵ_j_* are approximately i.i.d. 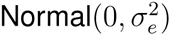. When a correlation between the adjusted trait and the adjusted genotype for SNP *i* is carried out, then the residual variation is based on the joint distribution of 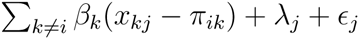 for *j* = 1, …, *n*.

Let us denote 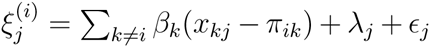. Since Var(*x_ij_* − *π_ij_*) = 2*π_ij_* (1 − *π_ij_*) and Var(*x_kj_* − *π_kj_*) = 2*π_kj_* (1 − *π_kj_*), it follows that (*x_ij_* − *π_ij_*) and (*x_kj_* − *π_kj_*) for *i, k* = 1, …, *m* and *j* = 1, …, *n* still suffer from confounding due to structure through their variances. Therefore, the implicit assumption made by the PCA approach that the 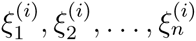 are independent and identically distributed in the above model is violated. This is our interpretation of why the PCA approach shows poor performance in adjusting for structure under our quantitative trait simulations. Astle and Balding (2009) [7] make further mathematical characterizations of the relationship between the implicit models in the PCA and LMM approaches, which we also found to be helpful.

Interestingly, when considering the binary trait model (2), the Bernoulli distributed trait does not involve a mean and variance term as in the Normal distributed quantitative trait. It may be the case that this difference contributes to explaining why the PCA approach shows similar behavior to the GCAT and LMM approaches for binary traits (see ref. [7]). Specifically, the PCA approach appears to perform reasonably well in adjusting for structure for the binary trait simulations that we considered.

#### 9 Software Implementation

The proposed method has been implemented in open source software, which is available at https://github.com/StoreyLab/gcat.

### Supplementary Figures

**Supplementary Figure 1:**
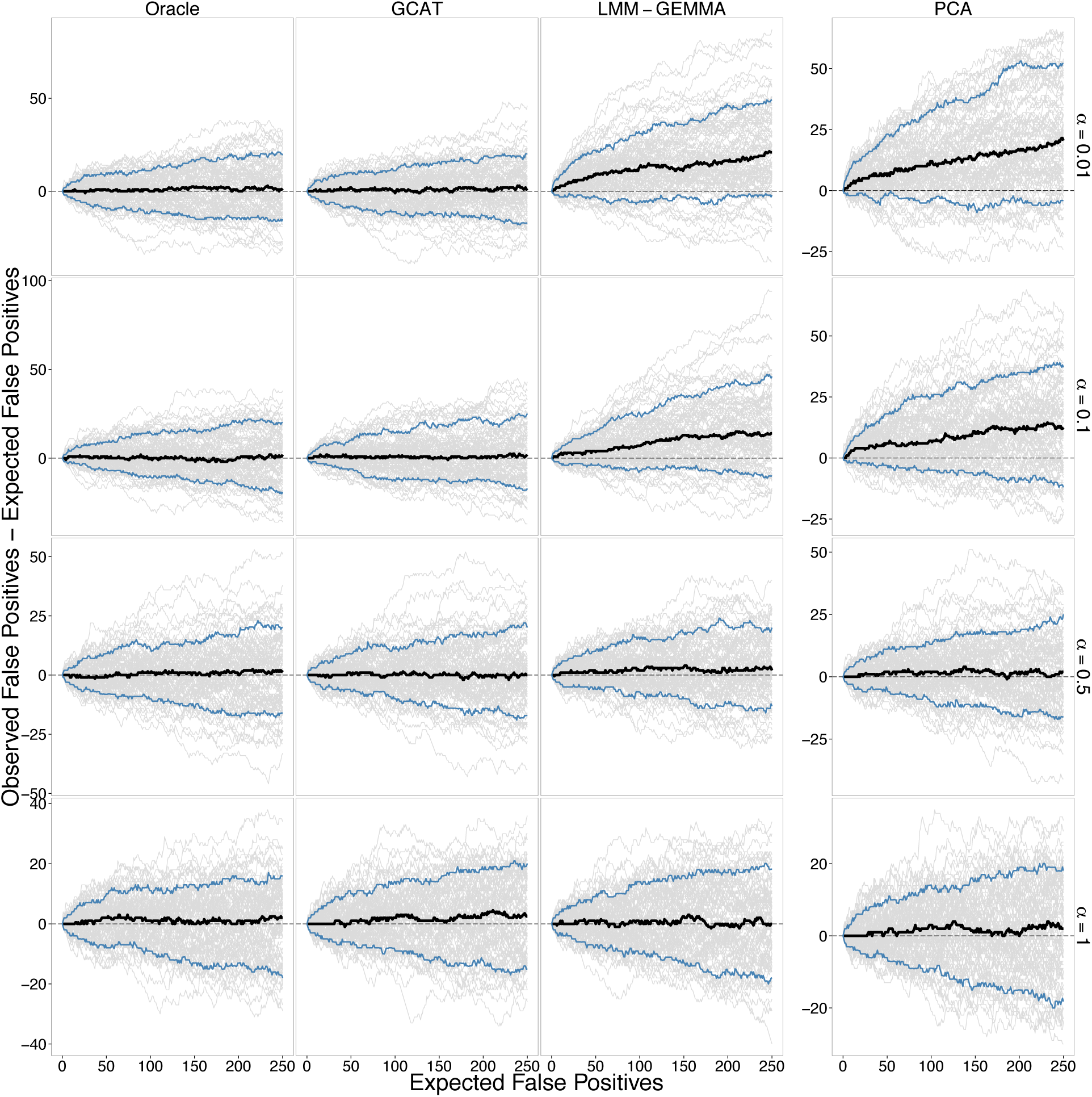
Performance of association tests on 100 simulated studies from the PSD model of structure for various *α* comparing the Oracle, GCAT (proposed), LMM-GEMMA, and PCA tests. The variance contributions to the trait are genetic=5%, environmental=5%, and noise=90%. The remaining details are equivalent to Figure 2.

**Supplementary Figure 2:**
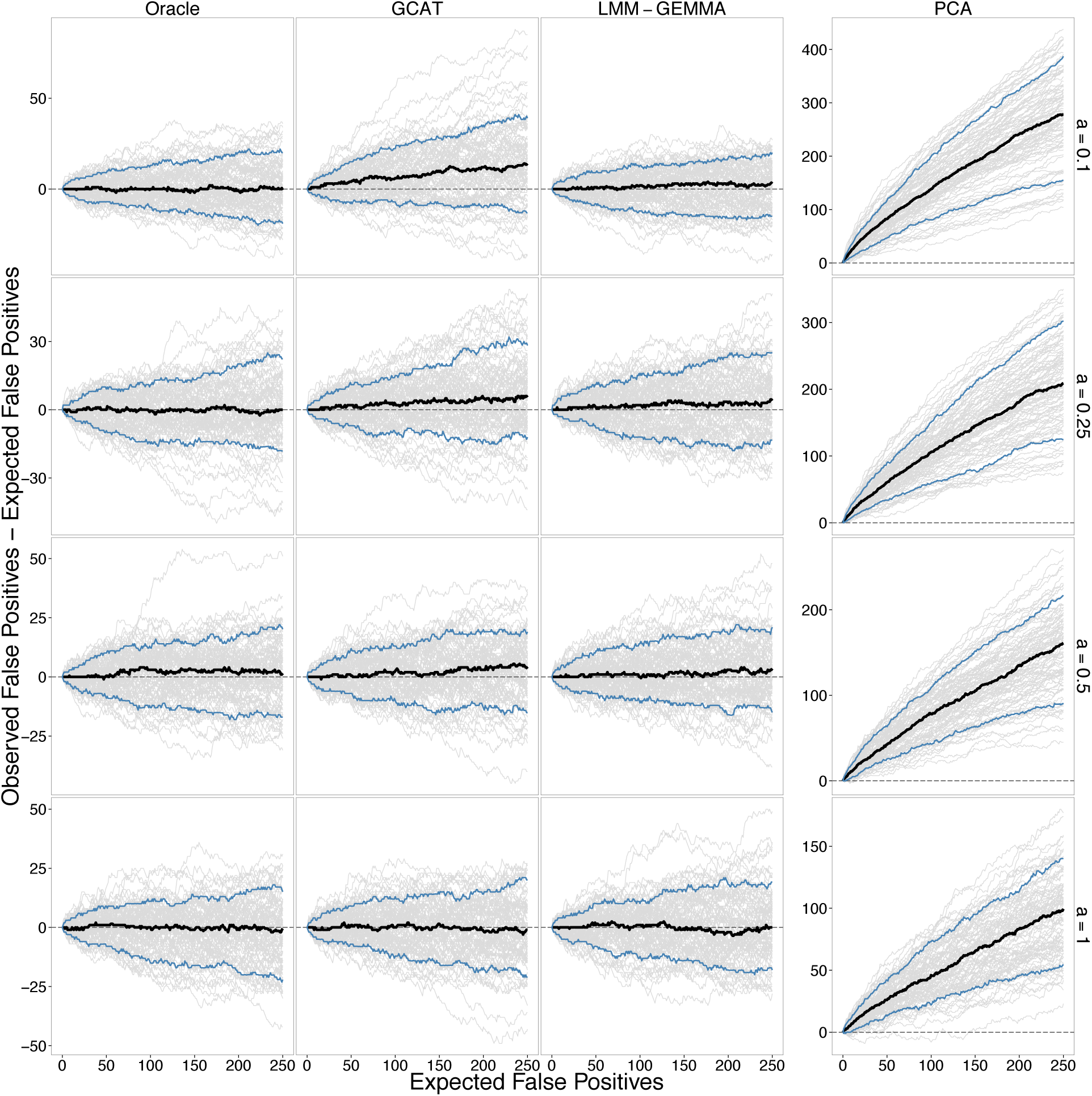
Performance of association tests on 100 simulated studies from the spatial model of structure for various *a* comparing the Oracle, GCAT (proposed), LMM-GEMMA, and PCA tests. The variance contributions to the trait are genetic=5%, environmental=5%, and noise=90%. The remaining details are equivalent to Figure 2.

**Supplementary Figure 3:**
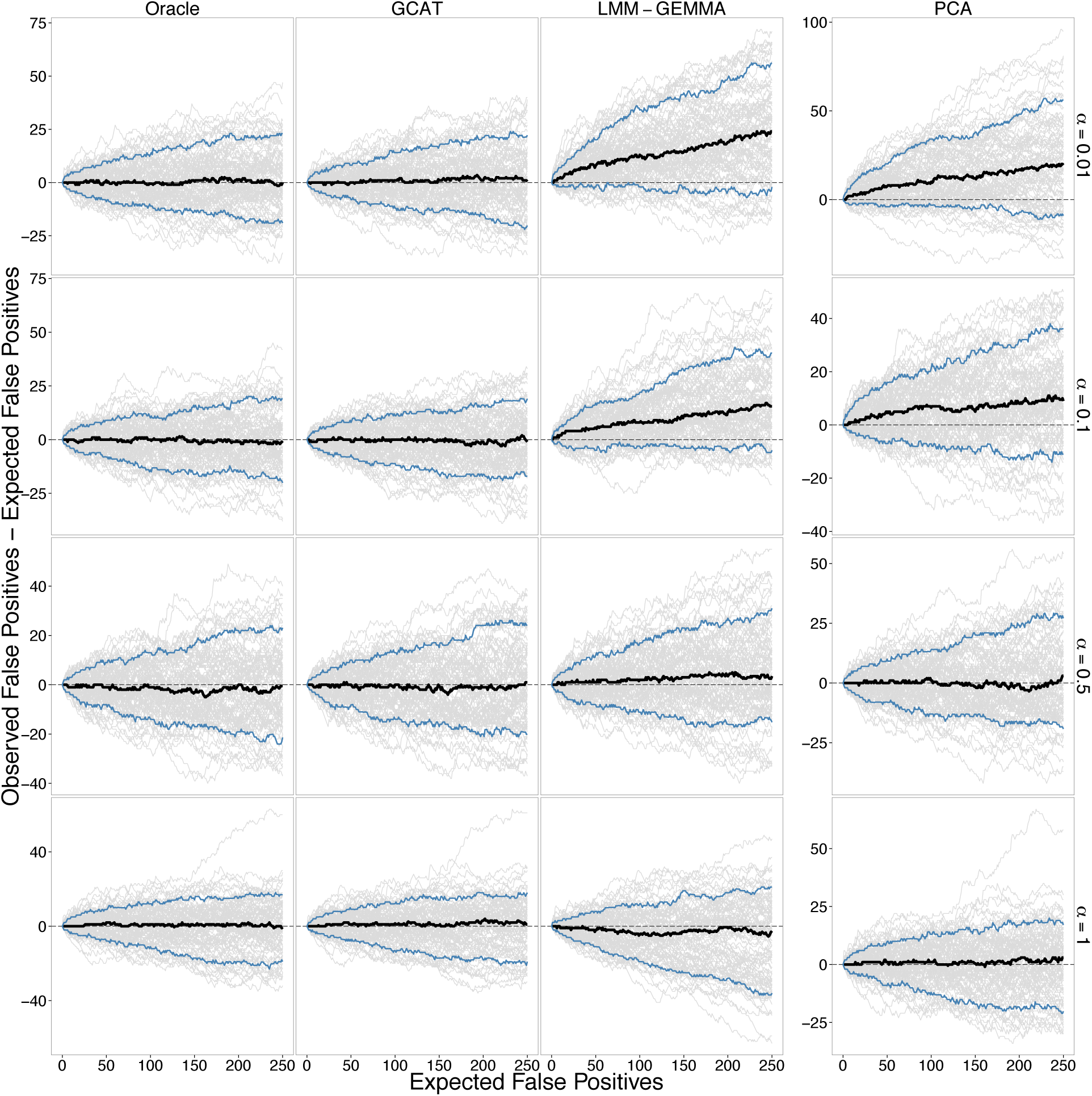
Performance of association tests on 100 simulated studies from the PSD model of structure for various *α* comparing the Oracle, GCAT (proposed), LMM-GEMMA, and PCA tests. The variance contributions to the trait are genetic=10%, environmental=0%, and noise=90%. The remaining details are equivalent to Figure 2.

**Supplementary Figure 4:**
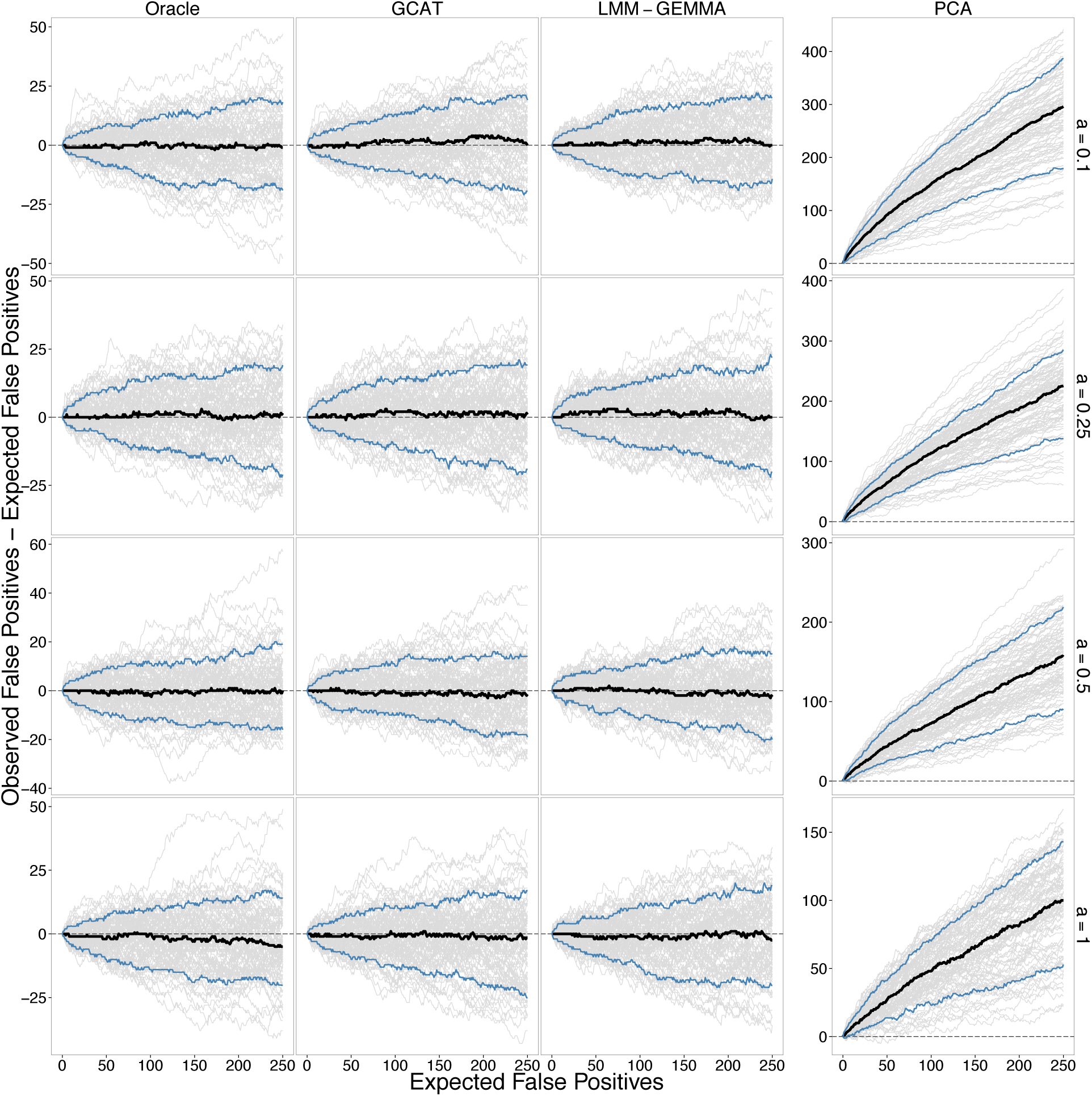
Performance of association tests on 100 simulated studies from the spatial model of structure for various *a* comparing the Oracle, GCAT (proposed), LMM-GEMMA, and PCA tests. The variance contributions to the trait are genetic=10%, environmental=0%, and noise=90%. The remaining details are equivalent to Figure 2.

**Supplementary Figure 5:**
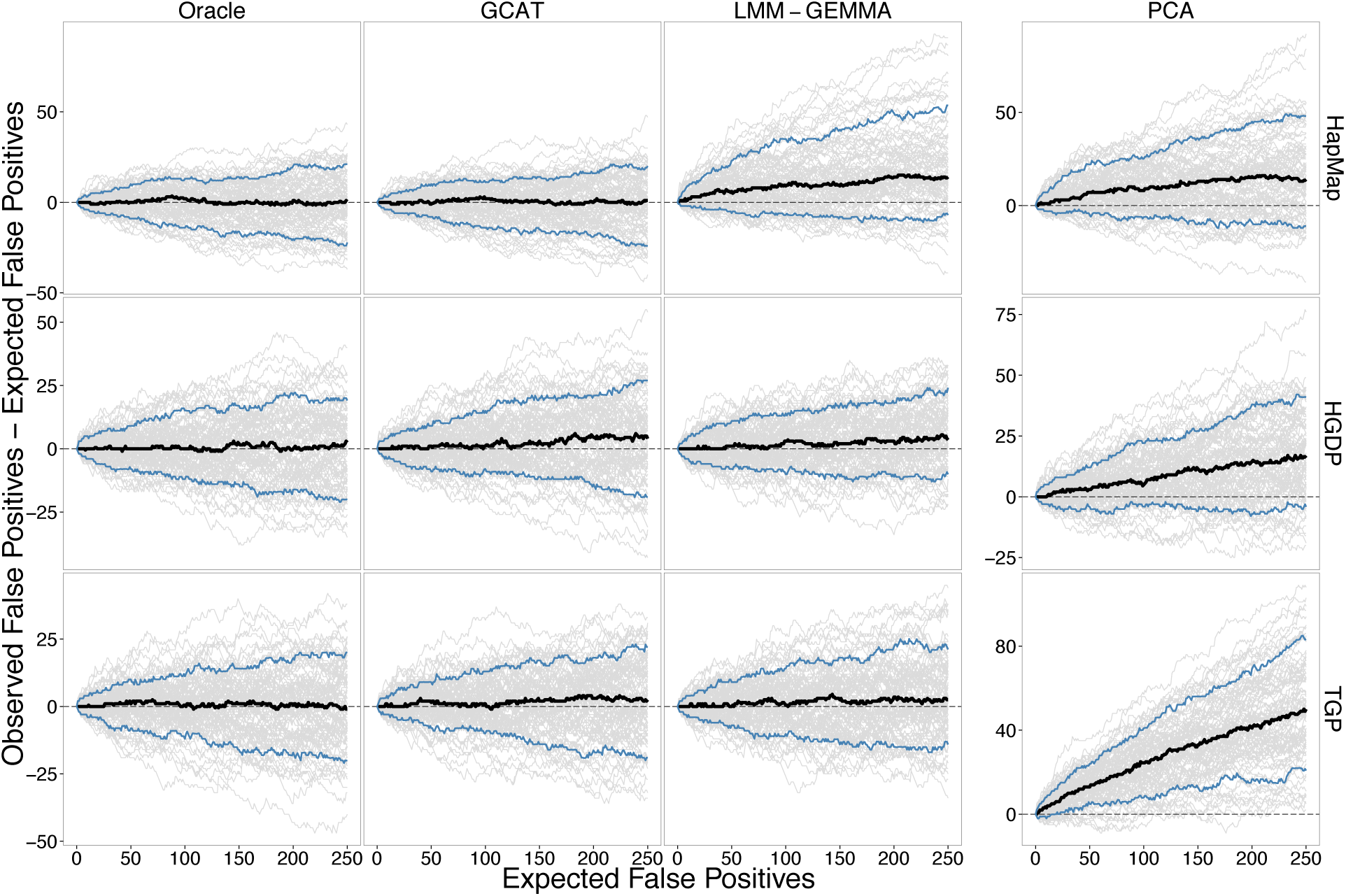
Performance of association tests on 100 simulated studies from the Balding-Nichols, HGDP, and TGP simulation scenarios comparing the Oracle, GCAT (proposed), LMMGEMMA, and PCA tests. The variance contributions to the trait are genetic=10%, environmental=0%, and noise=90%. The remaining details are equivalent to Figure 2.

**Supplementary Figure 6:**
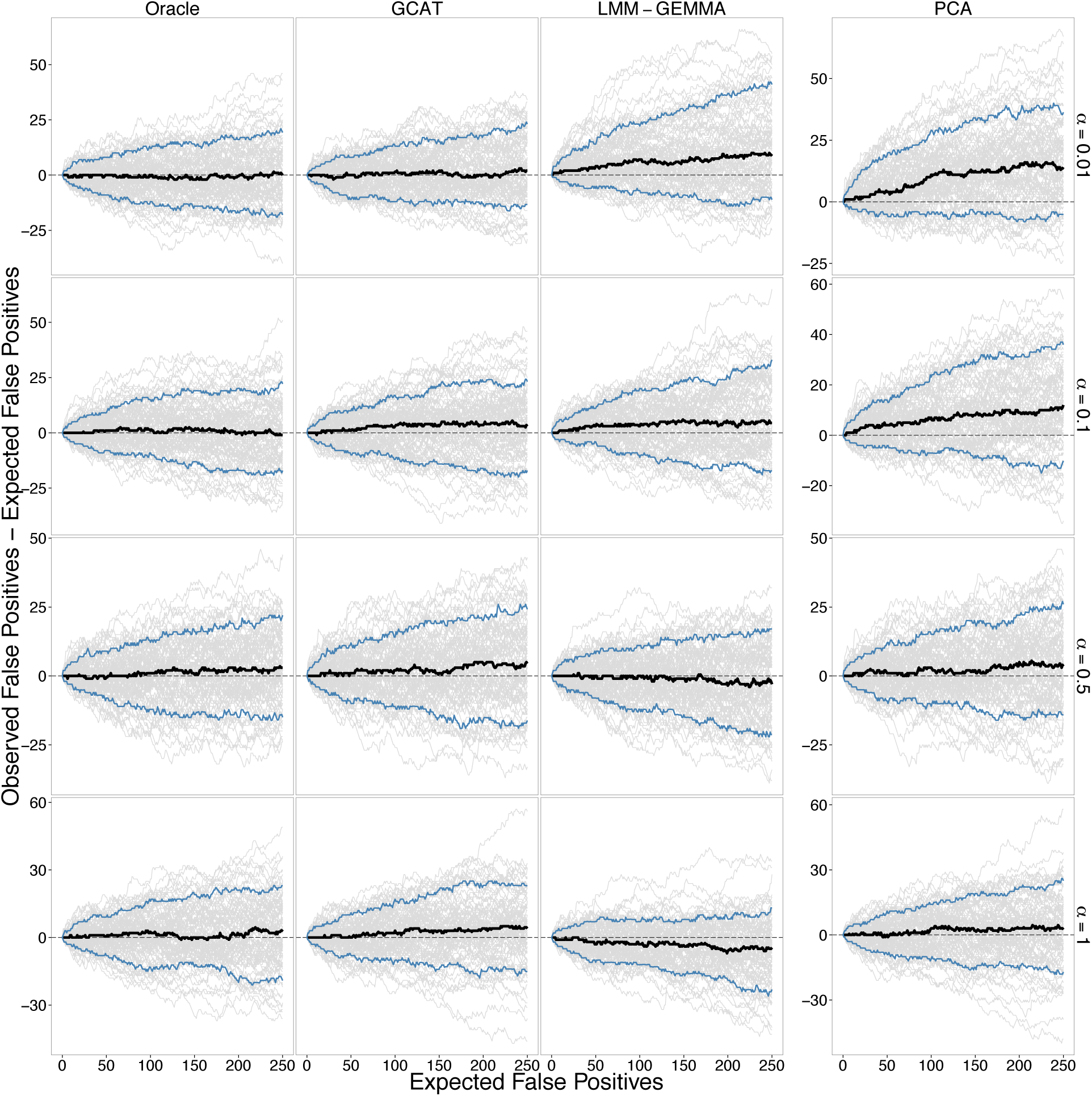
Performance of association tests on 100 simulated studies from the PSD model of structure for various *α* comparing the Oracle, GCAT (proposed), LMM-GEMMA, and PCA tests. The variance contributions to the trait are genetic=20%, environmental=10%, and noise=70%. The remaining details are equivalent to Figure 2.

**Supplementary Figure 7:**
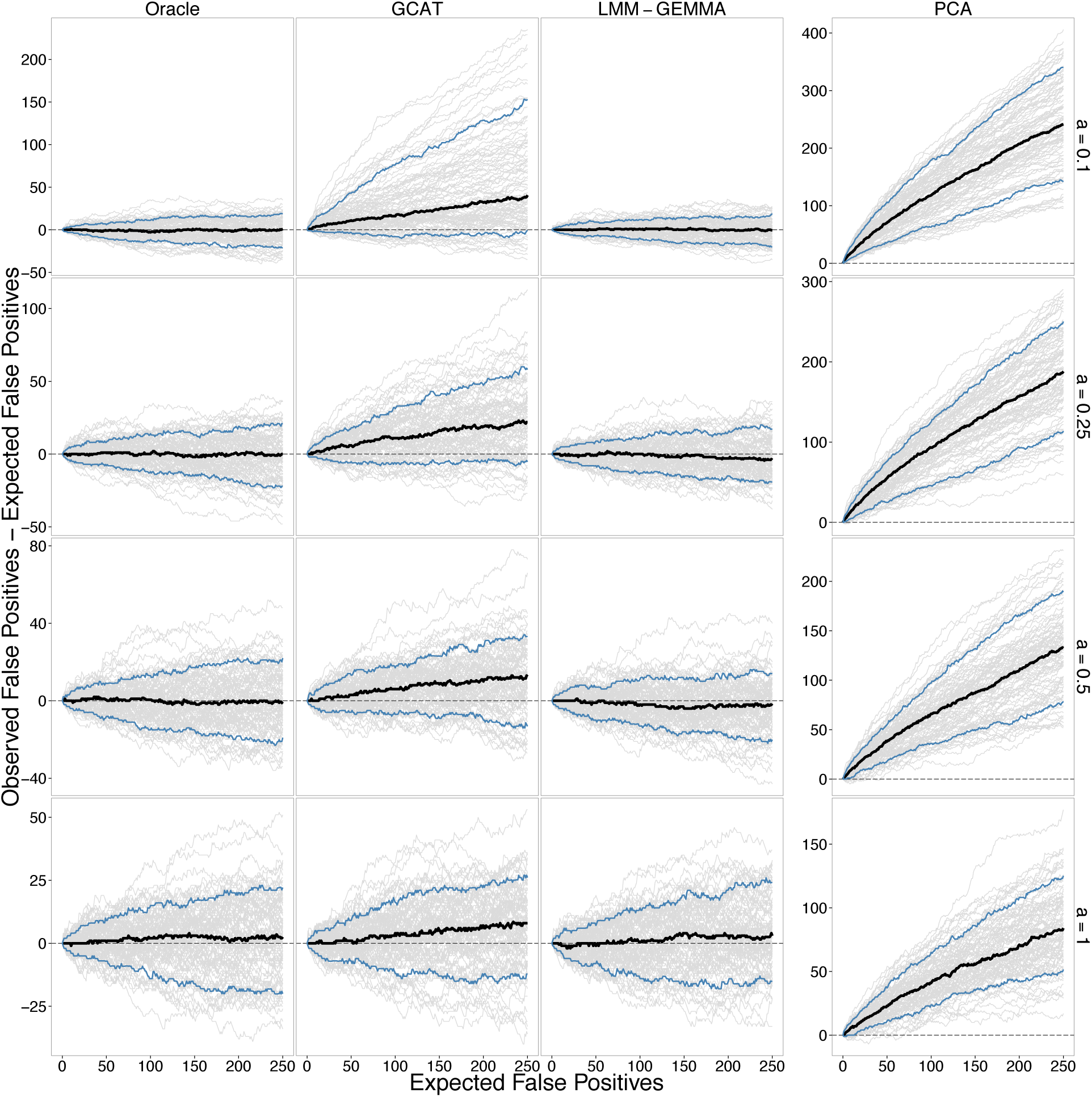
Performance of association tests on 100 simulated studies from the spatial model of structure for various *a* comparing the Oracle, GCAT (proposed), LMM-GEMMA, and PCA tests. The variance contributions to the trait are genetic=20%, environmental=10%, and noise=70%. The remaining details are equivalent to Figure 2. The difference in results between Oracle and GCAT is due to the fact that the *π_ij_* values are estimated in GCAT whereas the true *π_ij_* values are utilized in Oracle. In this particular simulation scenario, the error in *π_ij_* estimation results in a difference.

**Supplementary Figure 8:**
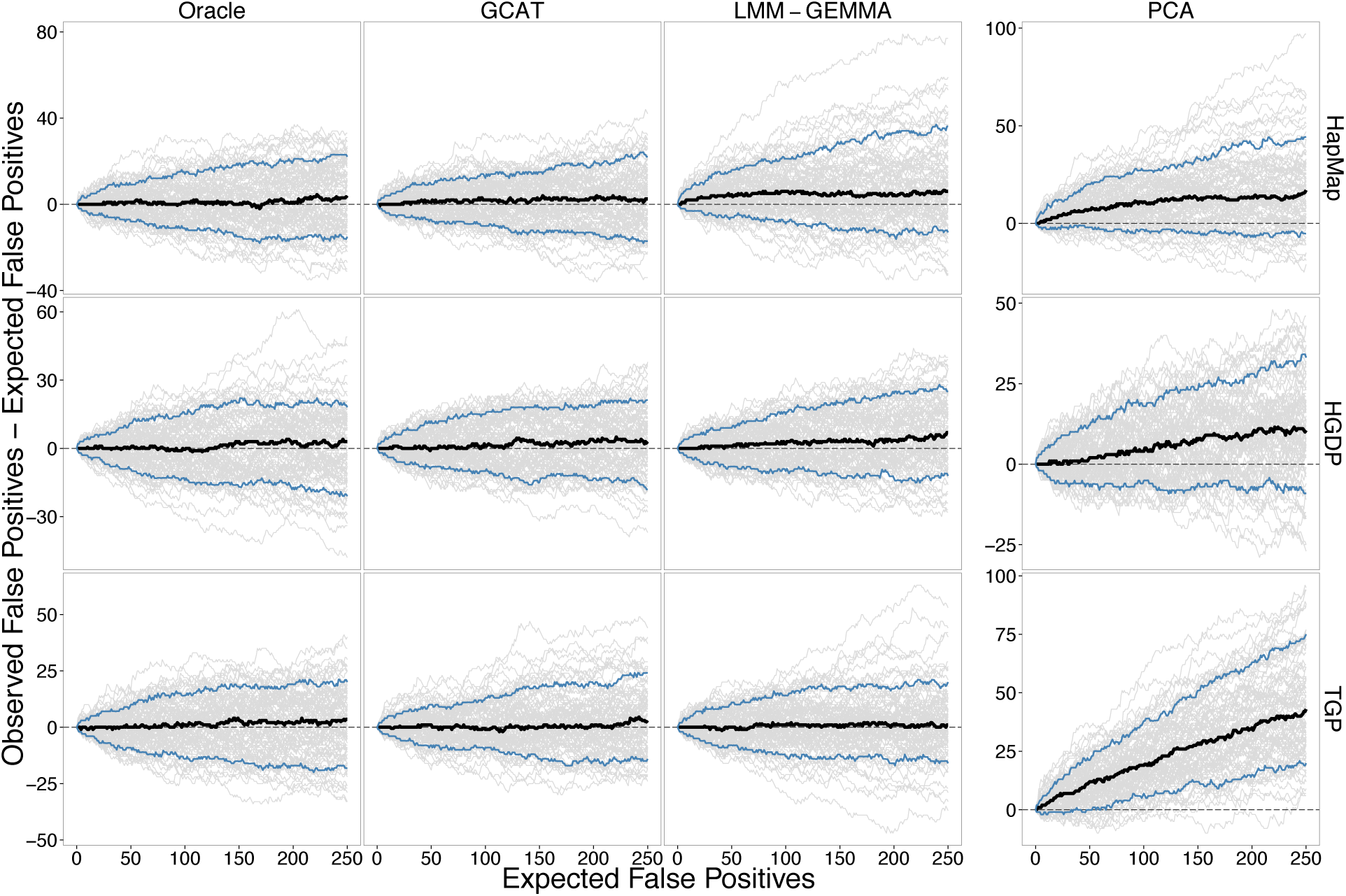
Performance of association tests on 100 simulated studies from the Balding-Nichols, HGDP, and TGP simulation scenarios comparing the Oracle, GCAT (proposed), LMMGEMMA, and PCA tests. The variance contributions to the trait are genetic=20%, environmental=10%, and noise=70%. The remaining details are equivalent to Figure 2.

**Supplementary Figure 9:**
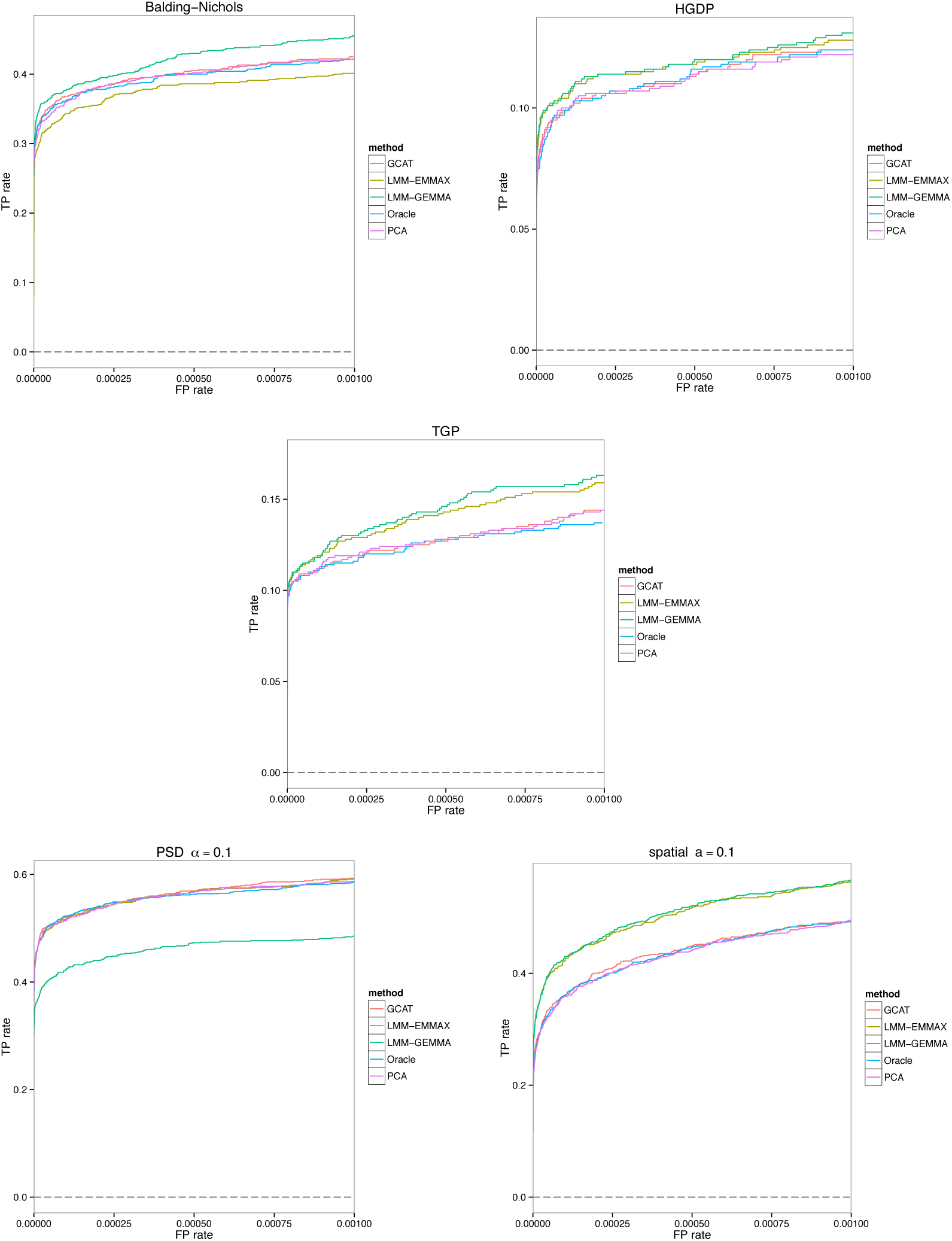
Statistical power of the Oracle, GCAT (proposed), PCA, and both LMM association tests. The results are for the simulated data sets shown in Figure 2. The quantitative traits are simulated from model (1) from Online Methods. The variance contributions to the trait are genetic=5%, environmental=5%, and noise=90%.

**Supplementary Figure 10:**
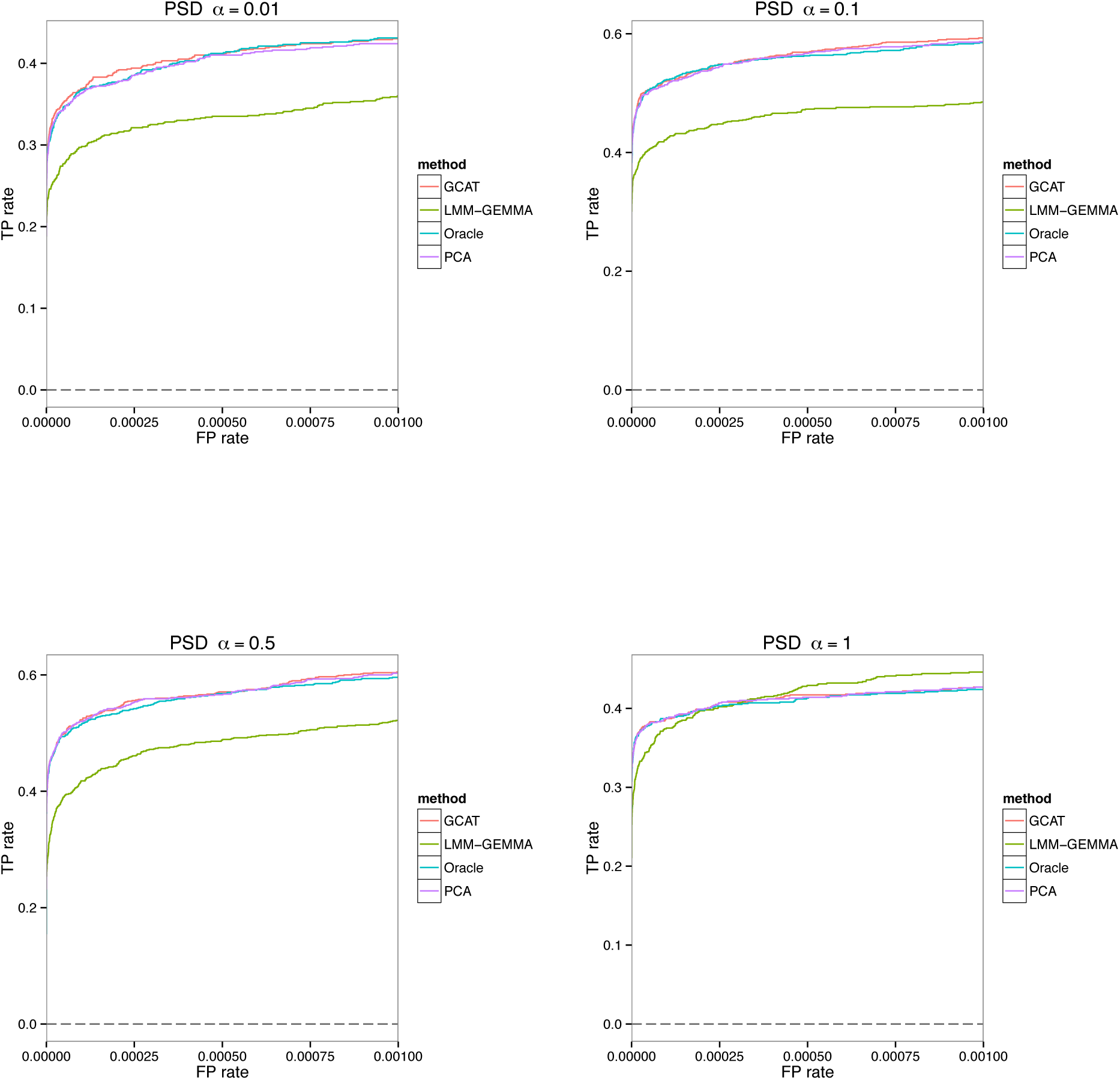
Power analysis for the simulation studies presented in Supplementary Figure 1.

**Supplementary Figure 11:**
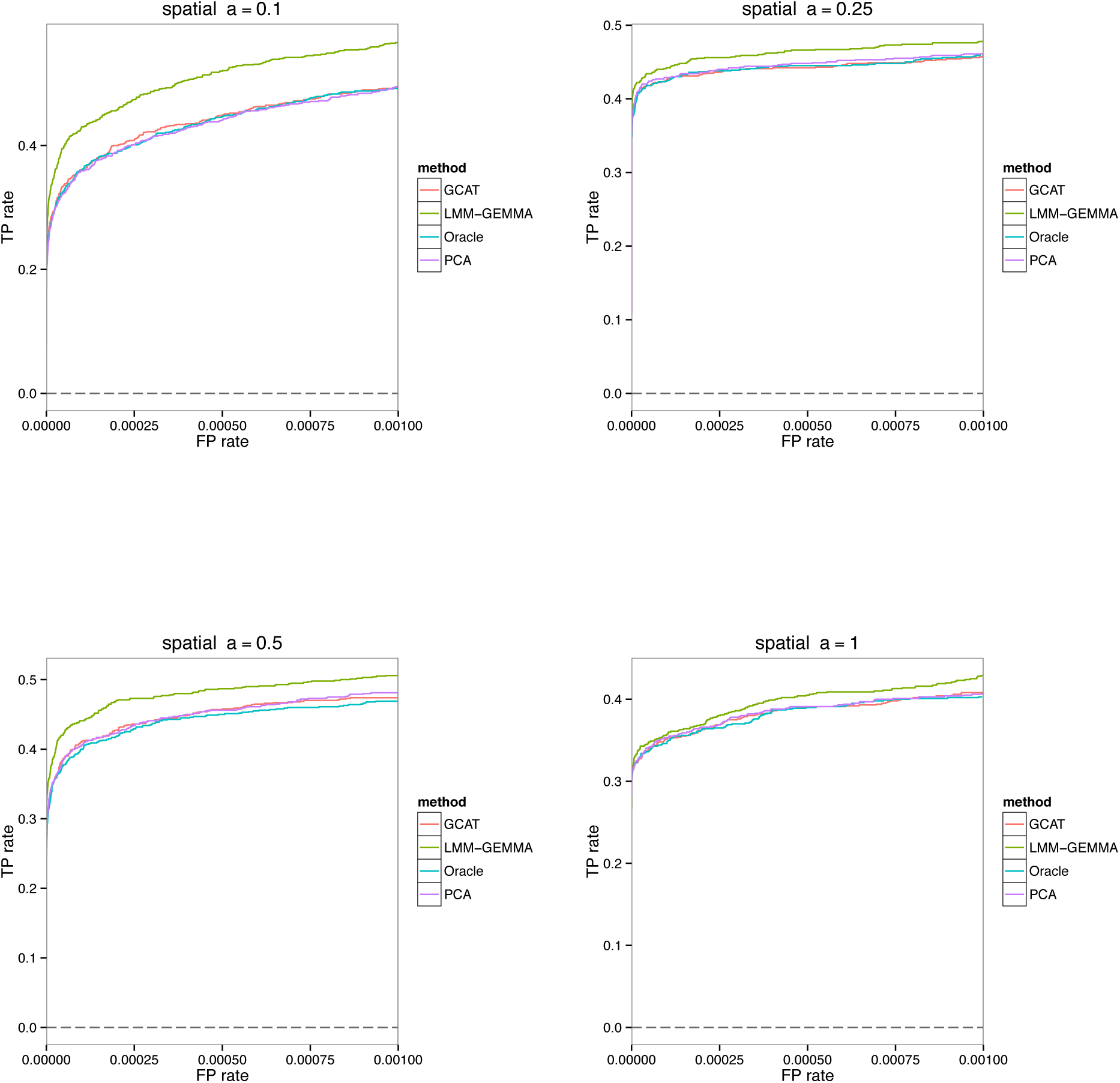
Power analysis for the simulation studies presented in Supplementary Figure 2.

**Supplementary Figure 12:**
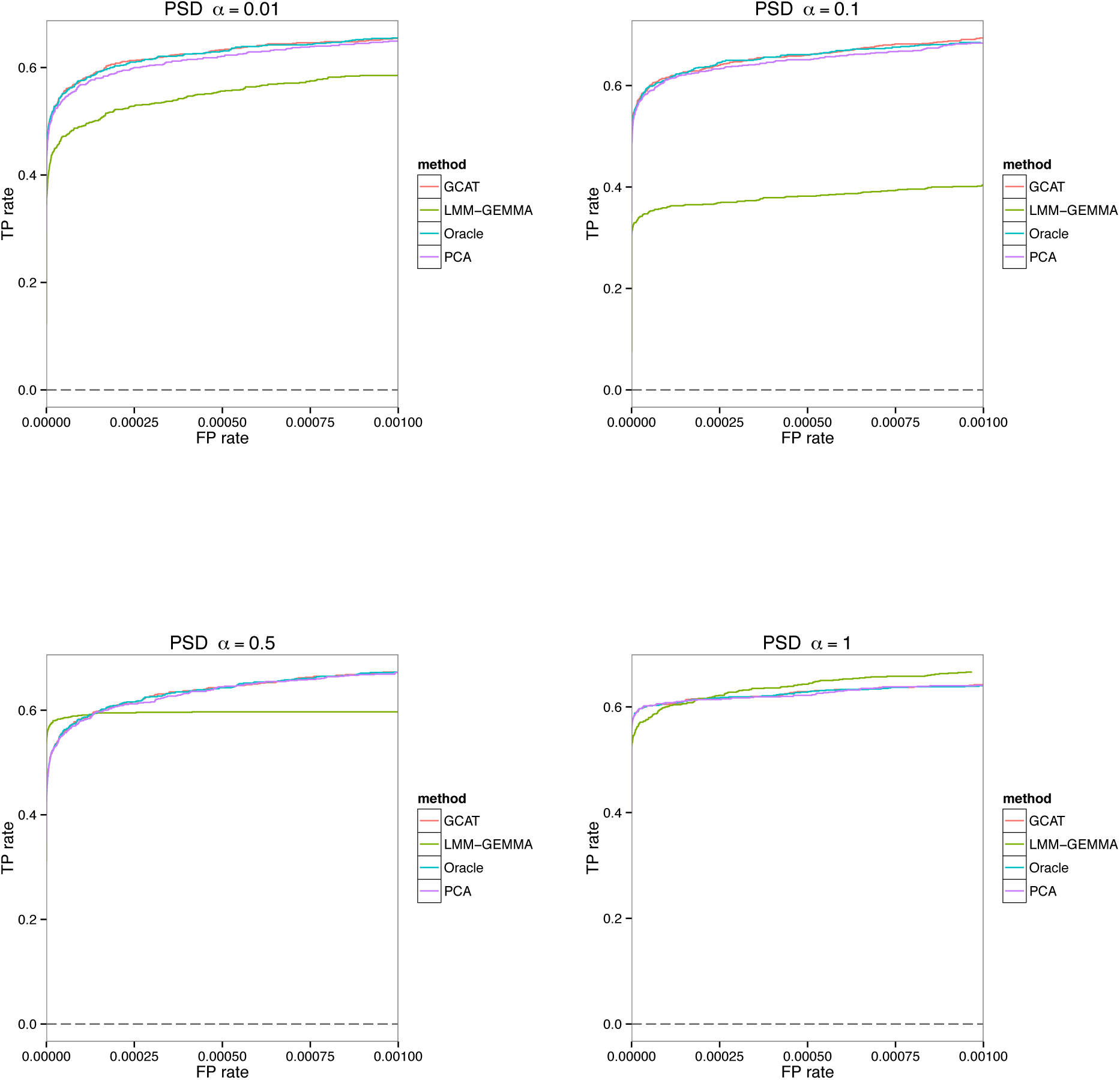
Power analysis for the simulation studies presented in Supplementary Figure 3.

**Supplementary Figure 13:**
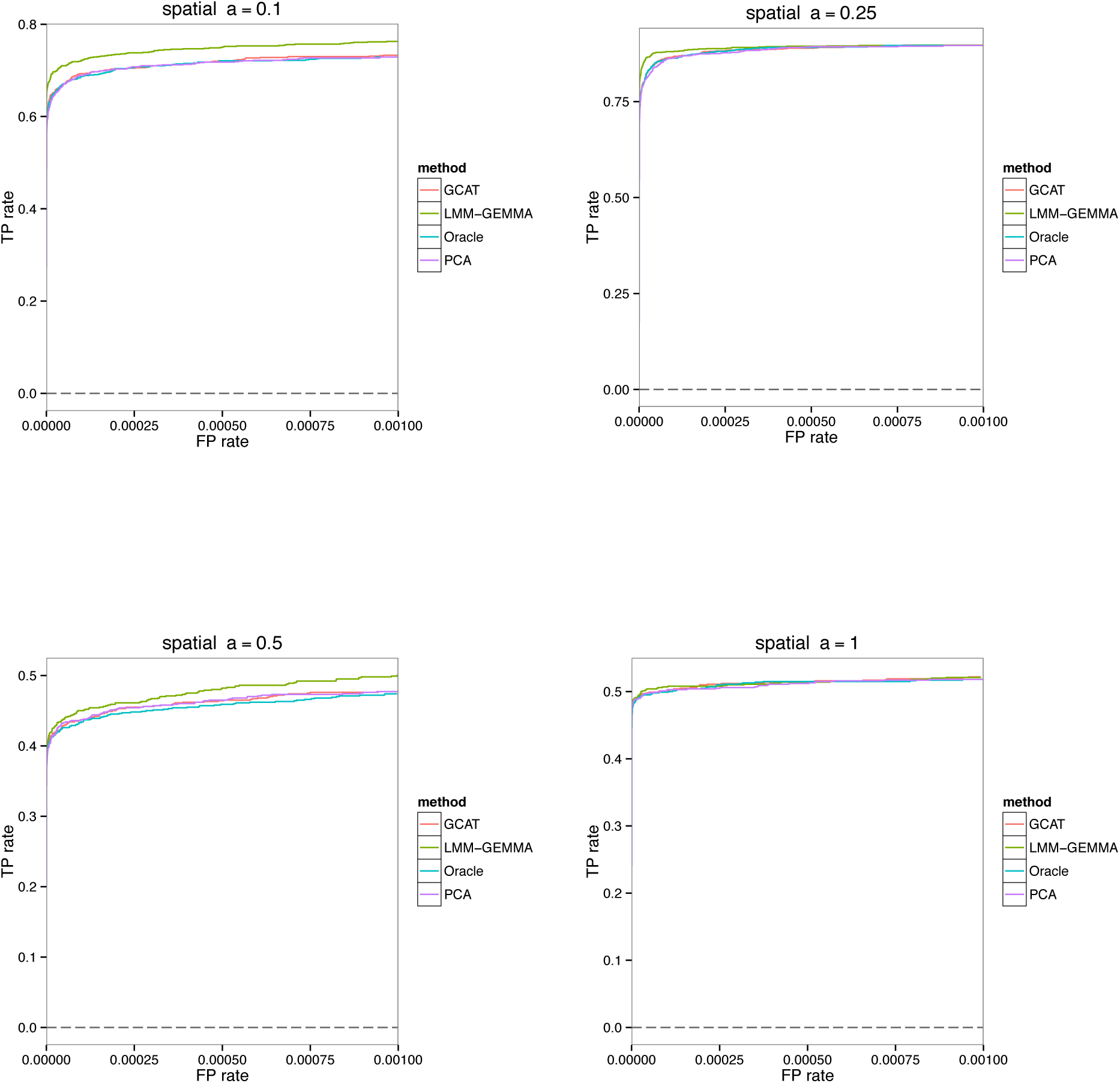
Power analysis for the simulation studies presented in Supplementary Figure 4.

**Supplementary Figure 14:**
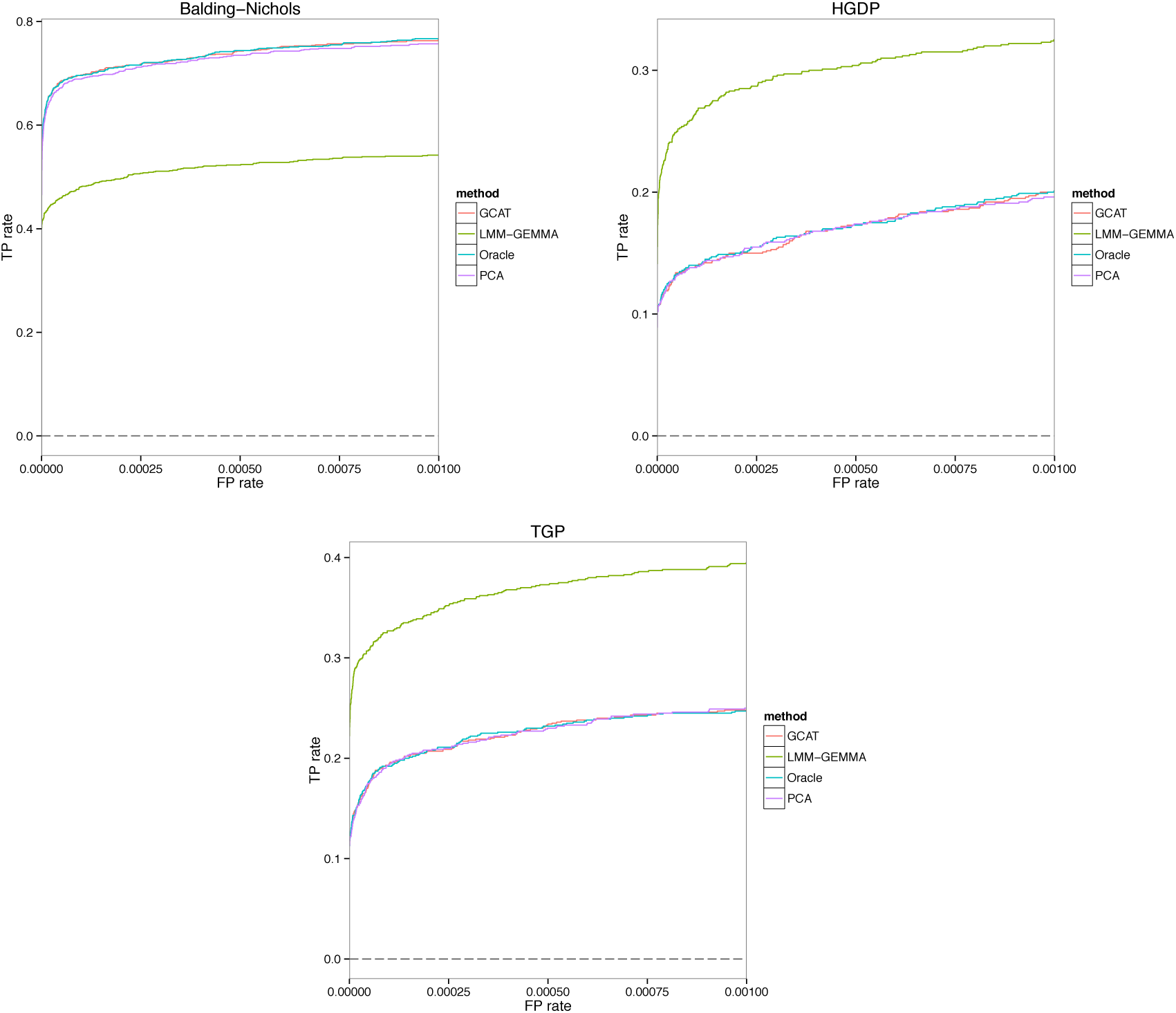
Power analysis for the simulation studies presented in Supplementary Figure 5.

**Supplementary Figure 15:**
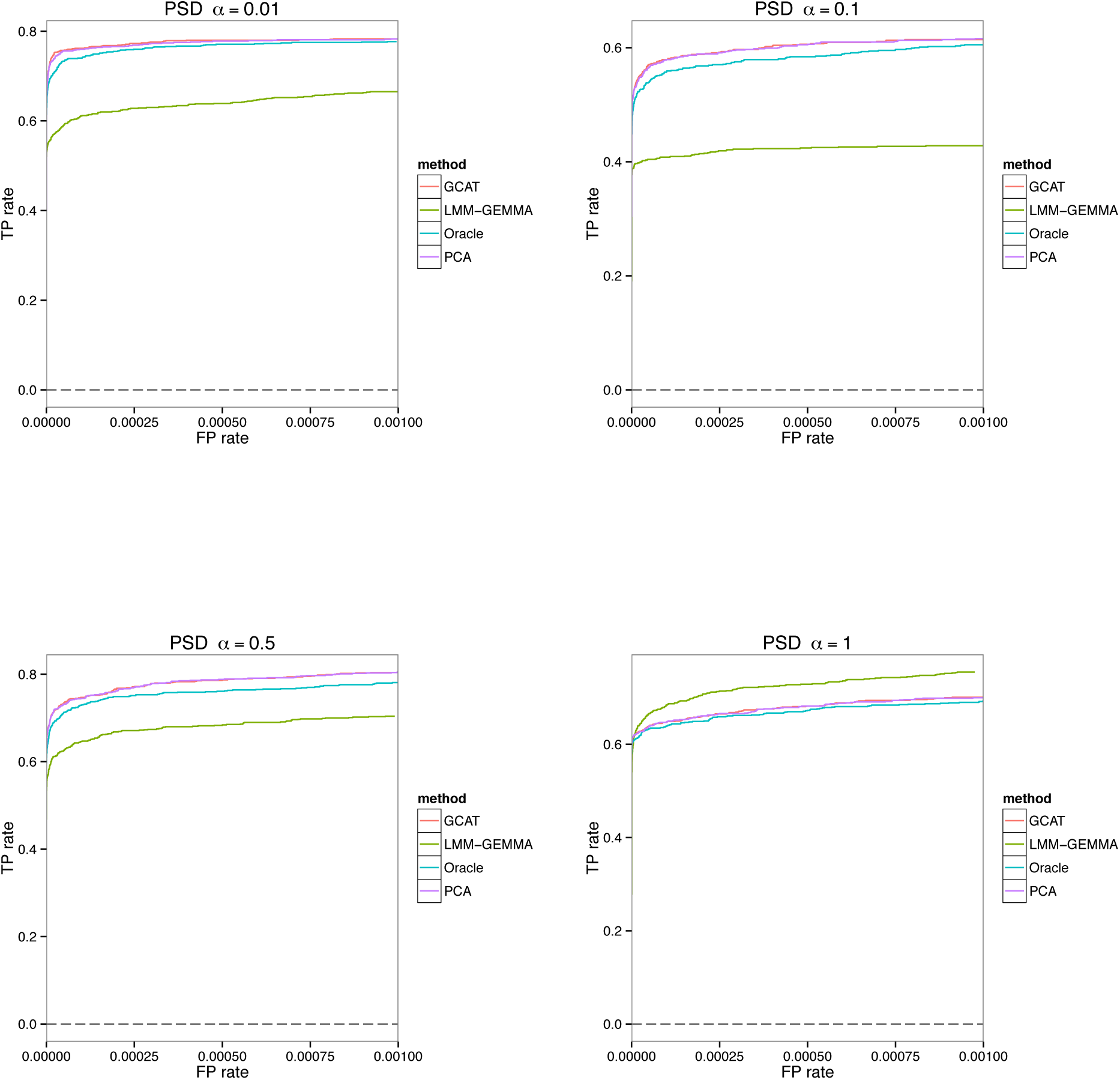
Power analysis for the simulation studies presented in Supplementary Figure 6.

**Supplementary Figure 16:**
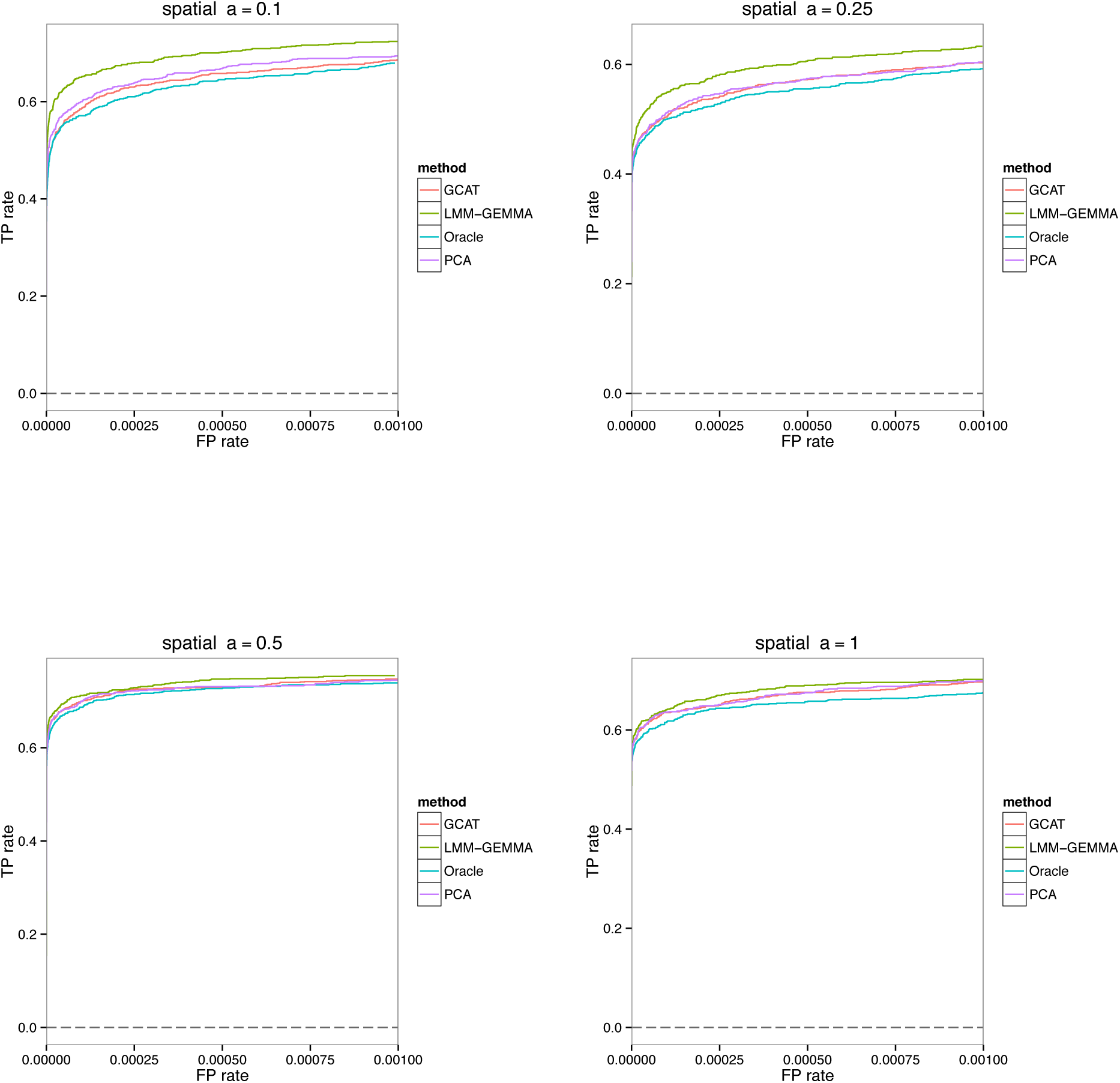
Power analysis for the simulation studies presented in Supplementary Figure 7.

**Supplementary Figure 17:**
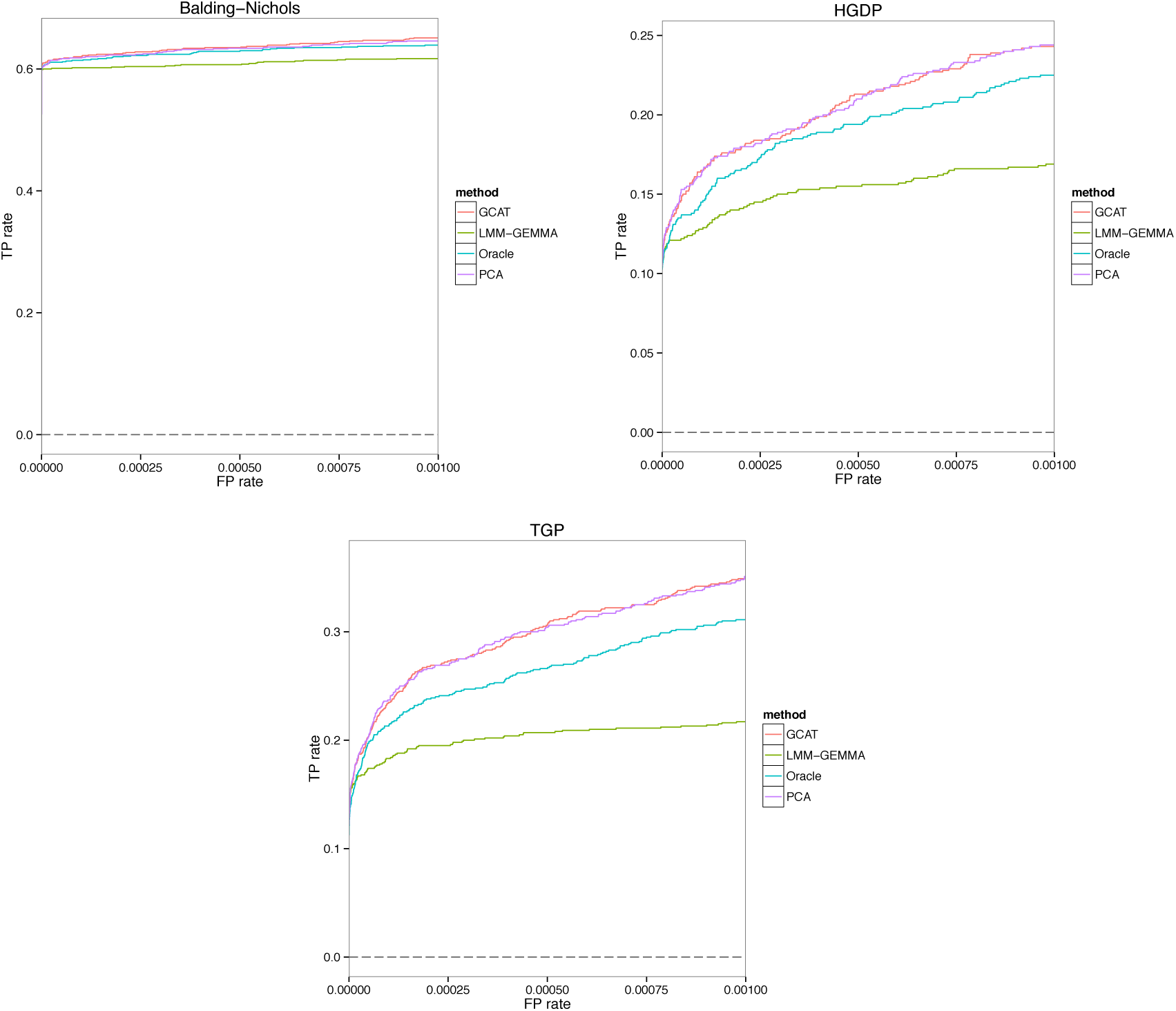
Power analysis for the simulation studies presented in Supplementary Figure 8.

**Supplementary Figure 18:**
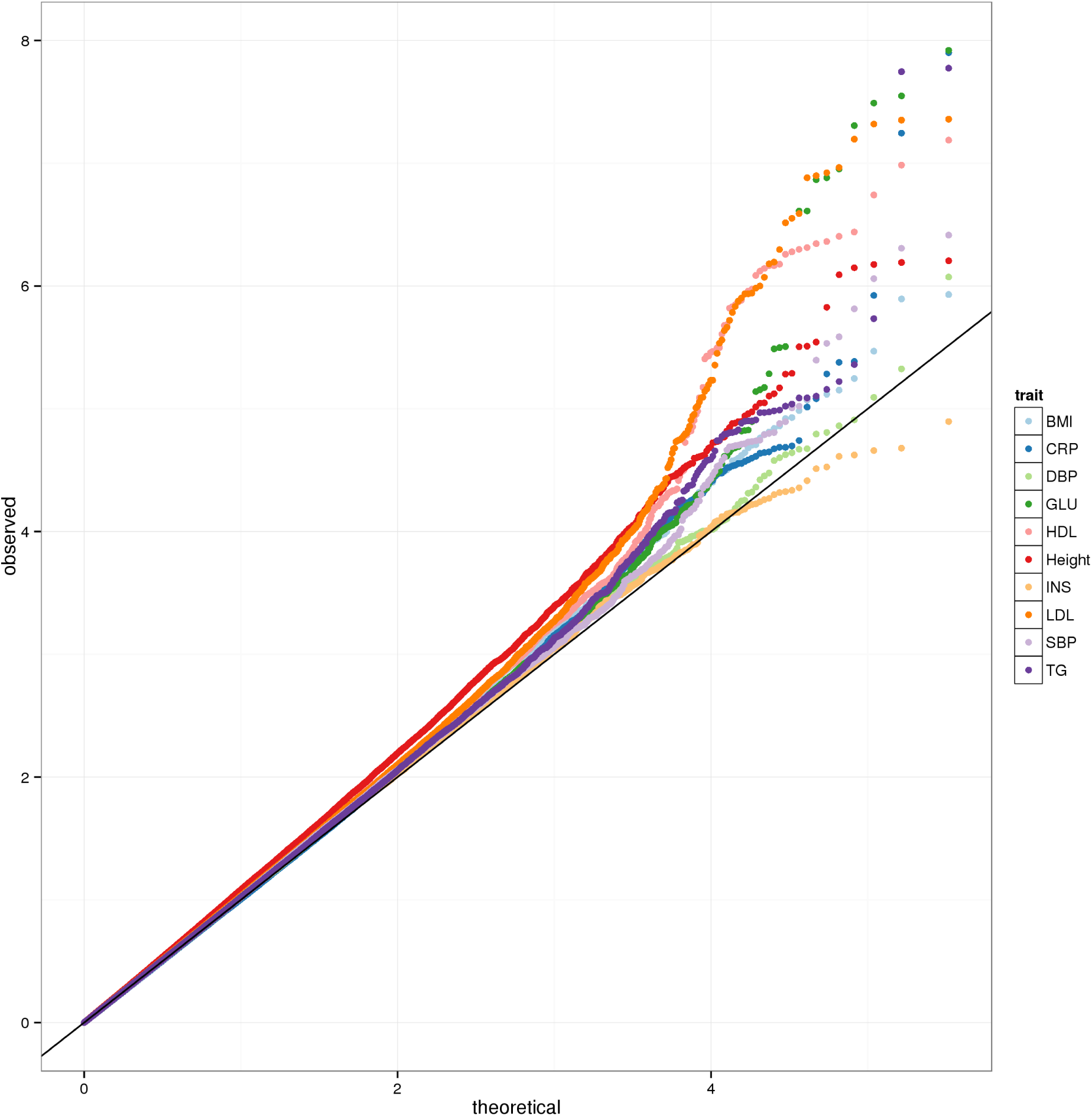
Theoretical versus observed quantiles of −log_10_(p-value) from the GCAT association tests on the Northern Finland Birth Cohort traits. The y-axis was truncated at p-value < 10^−8^; see Supplementary Table 1 for the smallest p-values for each trait.

### Supplementary Tables

**Supplementary Table 1:**
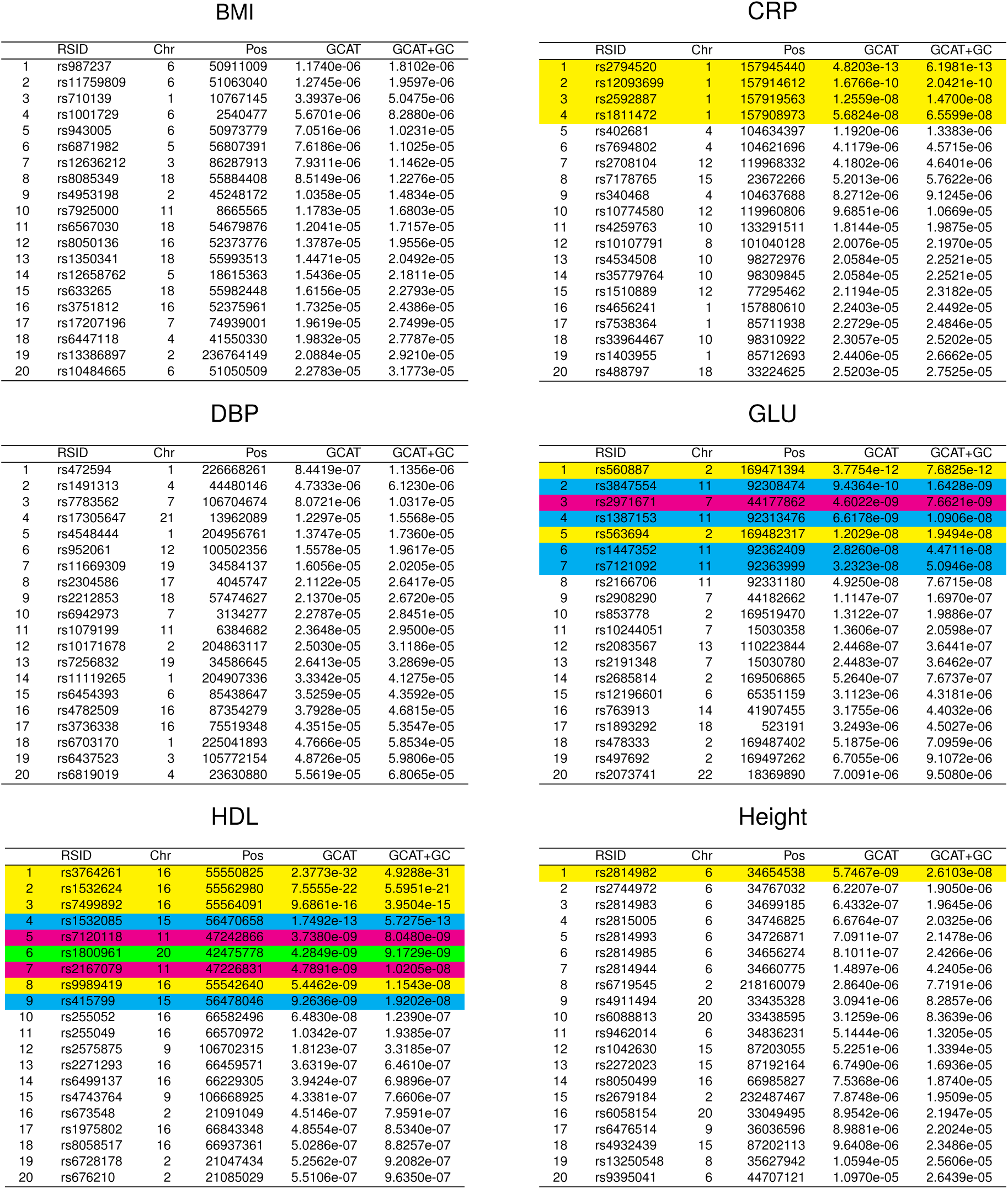
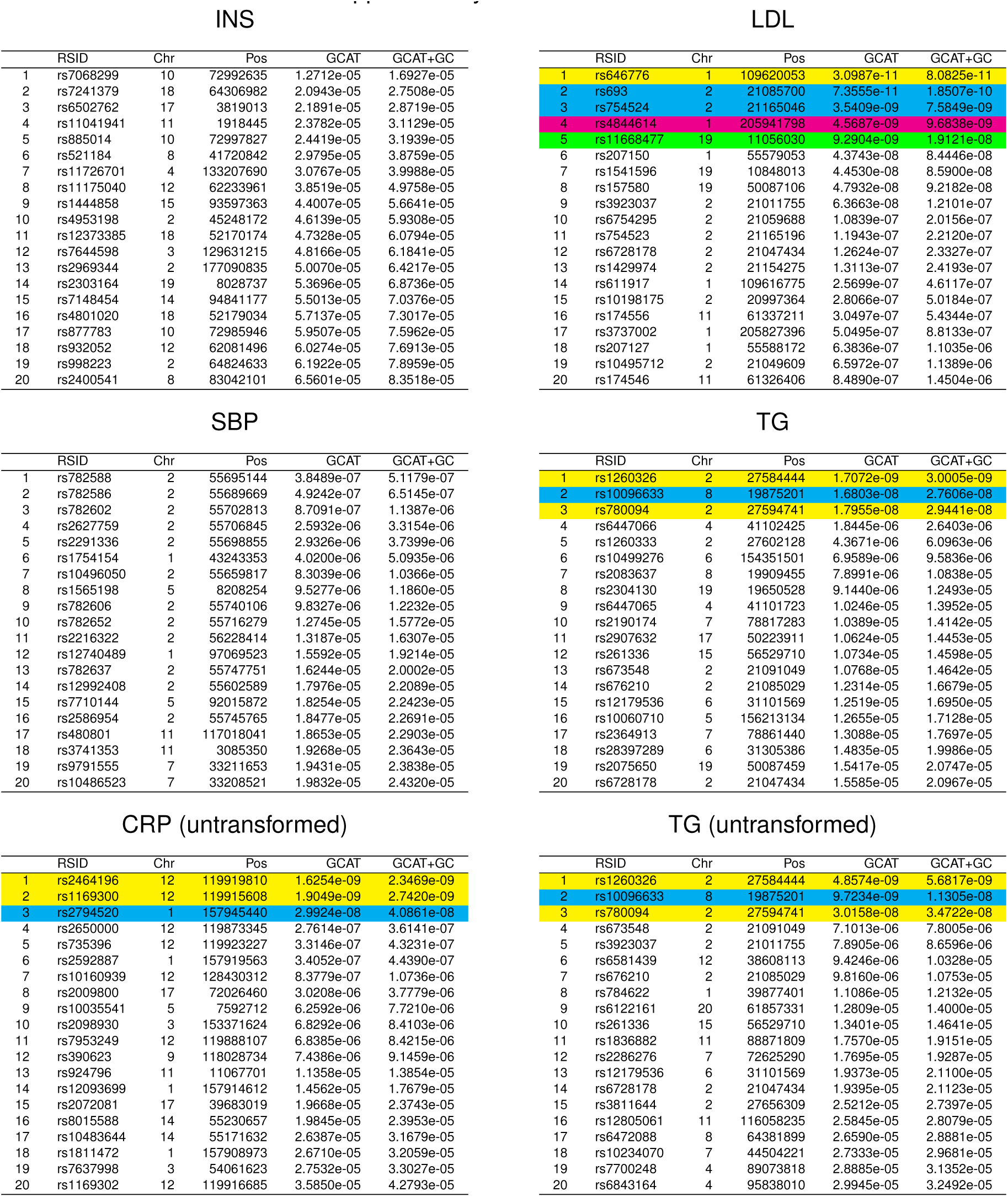
The top 20 most associated SNPs for each of the 10 traits considered in the Northern Finland Birth Cohort study. The GCAT p-value and GCAT+GC p-value (genomic control adjusted GCAT p-value) are shown for each SNP. SNPs that achieved GCAT+GC p-value < 7.2 × 10^−8^ are colored, and each locus for a given trait is given a different color.

**Supplementary Table 2:** The genomic control inflation factor (GCIF) was calculated for each trait in the association analysis of the Northern Finland Birth Cohort traits. The calculation was based on SNPs spaced at ~250kbp. The 95% Bonferroni adjusted simultaneous confidence interval under the assumption that the median statistic follows the theoretical null distribution is (0.9389, 1.0666). We calculated GCIF for the proposed statistics 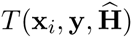 and 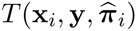 defined in the text.

| Trait | Abbreviation | $T(\mathbf{x}_i, \mathbf{y}, \hat{\mathbf{H}})$ | $T(\mathbf{x}_i, \mathbf{y}, \hat{\pi}_i)$ |
| --- | --- | --- | --- |
| Body Mass Index | BMI | 1.0633 | 1.0445 |
| C-reactive Protein | CRP | 1.0073 | 1.0050 |
| Diastolic blood pressure | DBP | 1.0487 | 1.0306 |
| Glucose | GLU | 1.0225 | 0.9886 |
| HDL Cholesterol | HDL | 1.0418 | 1.0206 |
| Height | Height | 1.0798 | 1.1017 |
| Insulin | INS | 1.0471 | 1.0636 |
| LDL Cholesterol | LDL | 1.0651 | 1.0264 |
| Systolic blood pressure | SBP | 1.0319 | 1.0336 |
| Triglycerides | TG | 1.0708 | 1.0327 |

